# Delaying the GABA shift indirectly affects membrane properties in the developing hippocampus

**DOI:** 10.1101/2023.02.13.528278

**Authors:** C. Peerboom, S. De Kater, N. Jonker, M. Rieter, T. Wijne, C.J. Wierenga

## Abstract

During the first two postnatal weeks intraneuronal chloride concentrations in rodents gradually decrease, causing a shift from depolarizing to hyperpolarizing γ-aminobutyric acid (GABA) responses. The postnatal GABA shift is delayed in rodent models for neurodevelopmental disorders and in human patients, but the impact of a delayed GABA shift on the developing brain remain obscure. Here we examine the direct and indirect consequences of a delayed postnatal GABA shift on network development in organotypic hippocampal cultures made from 6 to 7-day old mice by treating the cultures for one week with VU0463271, a specific inhibitor of the chloride exporter KCC2. We verified that VU treatment delayed the GABA shift and kept GABA signaling depolarizing until day in vitro (DIV) 9. We found that the structural and functional development of excitatory and inhibitory synapses at DIV9 was not affected after VU treatment. In line with previous studies, we observed that GABA signaling was already inhibitory in control and VU-treated postnatal slices. Surprisingly, fourteen days after the VU treatment had ended (DIV21), we observed an increased frequency of spontaneous inhibitory post-synaptic currents in CA1 pyramidal cells, while excitatory currents were not changed. Synapse numbers and release probability were unaffected. We found that dendrite-targeting interneurons in the *stratum Radiatum* had an elevated resting membrane potential, while pyramidal cells were less excitable compared to control slices. Our results show that depolarizing GABA signaling does not promote synapse formation after P7, and suggest that postnatal intracellular chloride levels indirectly affect membrane properties in a cell-specific manner.

**Significance Statement:** During brain development the action of neurotransmitter GABA shifts from depolarizing to hyperpolarizing. This shift is a thought to play a critical role in synapse formation. A delayed shift is common in rodent models for neurodevelopmental disorders and in human patients, but its consequences for synaptic development remain obscure. Here, we delayed the GABA shift by one week in organotypic hippocampal cultures and carefully examined the consequences for circuit development. We find that delaying the shift has no direct effects on synaptic development, but instead leads to indirect, cell type-specific changes in membrane properties. Our data call for careful assessment of alterations in cellular excitability in neurodevelopmental disorders.

## Introduction

During postnatal development of rodents intracellular chloride concentrations in neurons gradually decrease. Since GABA_A_ receptors mainly conduct chloride, the developmental decrease in intracellular chloride causes the reversal potential for GABA currents to gradually drop below resting membrane potential. As a result, the GABAergic driving force shifts from depolarizing in immature neurons to hyperpolarizing in mature neurons (Rivera et al., 1999; Hübner et al., 2001; Tyzio et al., 2007; Romo-Parra et al., 2008; Kirmse et al., 2015; Tsukahara et al., 2015; Sulis Sato et al., 2017; Murata and Colonnese, 2020). This GABA shift is a key event during development and is suggested to play a critical role in the formation and maturation of synapses (Leinekugel et al., 1997; Akerman and Cline, 2006; Wang and Kriegstein, 2008, 2011; Chancey et al., 2013; van Rheede et al., 2015; Oh et al., 2016).

The shift in postnatal chloride levels is the direct result of an increased function of the chloride exporter KCC2 (K-Cl cotransporter 2) relative to the chloride importer NKCC1 (Na-K-2Cl cotransporter 1) (Rivera et al., 1999; Gulyás et al., 2001; Yamada et al., 2004; Dzhala et al., 2005; Otsu et al., 2020). In humans, the shift in chloride transporter expression occurs during the first year after birth (Dzhala et al., 2005; Sedmak et al., 2016), but in some patients with neurodevelopmental disorders (NDDs) the GABA shift appear delayed and alterations in the expression of both NKCC1 and KCC2 have been widely reported (Talos et al., 2012; Duarte et al., 2013; Merner et al., 2015; Ruffolo et al., 2016; Birey et al., 2022). In rodents, the GABA shift occurs during the first two postnatal weeks (Rivera et al., 1999; Stein et al., 2004; Tyzio et al., 2007; Romo-Parra et al., 2008; Kirmse et al., 2015; Sulis Sato et al., 2017; Murata and Colonnese, 2020). In many animal models for NDDs a delayed GABA shift and alterations in chloride cotransporter expression have been reported (He et al., 2014; Tyzio et al., 2014; Banerjee et al., 2016; Corradini et al., 2017; Fernandez et al., 2018; Roux et al., 2018; Lozovaya et al., 2019; Bertoni et al., 2021), suggesting this is a key feature of many NDDs. In NDD mouse models alterations in early spontaneous network activity are observed (Gonçalves et al., 2013; Cheyne et al., 2019), and coordination between excitatory and inhibitory transmission is impaired in adult mice (Gogolla et al., 2009; Antoine et al., 2019). These observations suggest that a delay in the GABA shift during early postnatal development disturbs synaptic connectivity and network development with consequences for adulthood (Meredith, 2015; Molnár et al., 2020). However, a direct link between the timing of the GABA shift and synaptic connectivity remains elusive.

Depolarizing, excitatory GABA signaling supports spontaneous oscillations in the immature rodent brain (Ben-Ari et al., 1989; Khazipov et al., 2004; Sipila et al., 2006; Rheims et al., 2008; Spoljaric et al., 2019) and can induce the formation and maturation of excitatory synapses until the first week after birth (Leinekugel et al., 1997; Wang and Kriegstein, 2008, 2011; van Rheede et al., 2015; Oh et al., 2016). After the GABA shift, the KCC2 protein takes over the synapse-promoting role, as it directly facilitates spine growth by promoting actin dynamics in spines and supports AMPA receptor insertion and confinement (Li et al., 2007; Gauvain et al., 2011; Fiumelli et al., 2013; Chevy et al., 2015; Puskarjov et al., 2017; Awad et al., 2018; Kesaf et al., 2020). The effect of depolarizing GABA on inhibitory synapses is less clear, as studies have reported both positive and negative effects (Chudotvorova et al., 2005; Akerman and Cline, 2006; Nakanishi et al., 2007; Wang and Kriegstein, 2008, 2011). It is important to note that the shift from depolarizing to hyperpolarizing GABA signaling does not necessarily go hand in hand with a shift from excitatory to inhibitory action. Depolarizing GABA can already have an inhibitory action and limit neuronal activity when GABA-mediated depolarization is subthreshold and opening of GABA_A_ receptors shunts excitatory inputs (Staley and Mody, 1992; Kirmse et al., 2015; Murata and Colonnese, 2020; Salmon et al., 2020). When GABA signaling becomes inhibitory, it restrains excitatory synapse formation (Kang, 2019; Salmon, 2020) and network oscillations disappear (Ben-Ari et al., 1989; Khazipov et al., 2004). Recent studies have suggested that depolarizing GABA is already inhibitory in the brain of newborn rodents after P3-7 (Kirmse et al., 2015; Valeeva et al., 2016; Murata and Colonnese, 2020). This would imply that the role of depolarizing GABA in promoting synapse formation is limited to the period before and up to the first week after birth.

Many NDD mouse models show a delay in the postnatal GABA shift, although the precise delay is variable. For instance, the GABA shift is delayed with three days in the cerebellum of mice exposed *in utero* to valproate (Roux et al., 2018), the delay is two to seven days in the hippocampus of Magel2 knock out mice (Bertoni et al., 2021), and six days in the cortex of fragile X mice (He et al., 2014). Longer delays have also been reported (Banerjee et al., 2016; Corradini et al., 2017; Fernandez et al., 2018; Lozovaya et al., 2019) Here, we carefully assessed the direct and indirect consequences of a delayed postnatal GABA shift on the development of synapses. We used hippocampal organotypic cultures, as in these slices the anatomy and the development of excitatory and inhibitory synapses is largely preserved compared to the *in vivo* situation (De Simoni, 2003). We treated the cultures with the KCC2 antagonist VU0463271 (VU) to block chloride extrusion for one week (from DIV2 to DIV8), while preserving the structural role of the KCC2 protein in spines (Li et al., 2007; Kesaf et al., 2020). VU treatment delayed the GABA shift without affecting expression levels of the chloride transporters. We found that excitatory and inhibitory synapses were not affected after one week of VU treatment. However, 14 days after the VU treatment had ended, we observed an increased frequency of spontaneous inhibitory postsynaptic currents (sIPSC), while excitatory postsynaptic currents (sEPSCs) were unaltered. We found that this was correlated with specific alterations in membrane excitability of pyramidal cells and interneurons. Our data suggest that delaying the GABA shift does not directly affect synaptic development, but rather leads to indirect cell-type specific changes in membrane properties.

## Materials and methods

### Animals

All animal experiments were performed in compliance with the guidelines for the welfare of experimental animals issued by the Federal Government of the Netherlands and were approved by the Dutch Central Committee Animal experiments (CCD), project AVD1080020173847 and project AVD1150020184927. In this study we used male and female transgenic mice: GAD65-GFP mice (López-Bendito et al., 2004) (bred as a heterozygous line in a C57BL/6JRj background), VGAT-Cre mice (JAX stock #028862) and SuperClomeleon (SClm) mice. The SClm mice (Herstel et al., 2022) are SuperClomeleon^lox/-^ mice (Rahmati et al., 2021), a gift from Kevin Staley (Massachusetts General Hospital, Boston, MA) crossed with CamKIIα^Cre/-^ mice (Tsien et al., 1996; Casanova et al., 2001). SClm mice express the chloride sensor SClm in up to 70% of the pyramidal neurons in the hippocampus (Casanova et al., 2001; Wang et al., 2013). VGAT-Cre mice express Cre recombinase expression in all inhibitory GABAergic neurons (Vong et al., 2011), and GAD65-GFP mice express GFP in ∼20% of GABAergic interneurons in the CA1 region of the hippocampus (Wierenga et al., 2010). As we did not detect any differences between slices from male and female mice, all data were pooled.

### Organotypic hippocampal culture preparation and VU treatment

Organotypic hippocampal cultures were made from P6-7 mice as described before (Hu et al., 2019; Herstel et al., 2022), based on Stoppini *et al*. (Stoppini et al., 1991). Mice were decapitated and their brain was rapidly placed in ice-cold Grey’s Balanced Salt Solution (GBSS; containing (in mM): 137 NaCl, 5 KCl, 1.5 CaCl_2_, 1 MgCl_2_, 0.3 MgSO_4_, 0.2 KH_2_PO_4_ and 0.85 Na_2_HPO_4_) with 25 mM glucose, 12.5 mM HEPES and 1 mM kynurenic acid (pH set at 7.2, osmolarity set at ∼320 mOsm, sterile filtered). 400 μm thick transverse hippocampal slices were cut with a McIlwain tissue chopper. Slices were placed on Millicell membrane inserts (Millipore) in wells containing 1 ml culture medium (consisting of 48% MEM, 25% HBSS, 25% horse serum, 25 mM glucose, and 12.5 mM HEPES, with an osmolarity of ∼325 mOsm and a pH of 7.3 – 7.4). Slices were stored in an incubator (35°C with 5% CO_2_). Culture medium was replaced by culture medium supplemented with 0.1% Dimethyl sulfoxide (DMSO, Sigma-Aldrich) or 1 μM VU VU0463271 (VU, Sigma-Aldrich, in 0.1 % DMSO) at DIV 1. In addition, a small drop of medium supplemented with DMSO or VU was carefully placed on top of the slices. In this way, cultures were treated 3 times per week until DIV8. From DIV8 onwards, cultures received normal culture medium 3 times per week. Experiments were performed at day in vitro 1-3 (DIV2), 7-8 (DIV8), 8-10 (DIV9) or 20-22 (DIV21). Please note that experiments at DIV9 were performed 1 to 55 hr after cessation of the VU treatment, but within groups there was no correlation between measurements and DIV.

### Electrophysiology and analysis

Organotypic cultures were transferred to a recording chamber and continuously perfused with carbonated (95% O_2_, 5% CO_2_) artificial cerebrospinal fluid (ACSF, in mM: 126 NaCl, 3 KCl, 2.5 CaCl_2_, 1.3 MgCl_2_, 26 NaHCO_3_, 1.25 NaH_2_PO_4_, 20 glucose; with an osmolarity of ∼310 mOsm) at a rate of approximately 1 ml/min. For acute treatments, ACSF containing 0.1% DMSO or 1 μM VU (dissolved in 0.1% DMSO) was bath applied for 5 minutes. Bath temperature was maintained at 29-32°C. Recordings were acquired using an MultiClamp 700B amplifier (Molecular Devices), filtered with a 3 kHz Bessel filter and stored using pClamp10 software.

Perforated patch clamp recordings were performed in CA1 pyramidal neurons. Recording pipettes with resistances of 2-4 MΩ were pulled from borosilicate glass capillaries (World Precision Instruments). The pipette tip was filled with gramicidin-free KCl solution (140 mM KCl and 10 mM HEPES, pH adjusted to 7.2 with KOH, and osmolarity 285 mOsm/l) and then backfilled with solution containing gramicidin (60 μg/ml, Sigma). Neurons were held at -65 mV and the access resistance of the perforated cells was monitored constantly before and during recordings. An access resistance of 50 MΩ was considered acceptable to start recording. Recordings were excluded if the resting membrane potential exceeded -50 mV or if the series resistance after the recording deviated more than 30% from its original value. GABAergic currents were induced by upon local application of 50 μM muscimol dissolved in HEPES-ACSF (containing in mM: 135 NaCl, 3 KCl, 2.5 CaCl_2_, 1.3 MgCl_2_, 1.25 Na_2_H_2_PO_4_, 20 Glucose, and 10 HEPES) every 30s using a Picospritzer II. Muscimol responses were recorded in the presence of 1 μM TTX (Abcam) at holding potentials between -100 mV and -30 mV in 10 mV steps. We plotted response amplitude as a function of holding potential and calculated the chloride reversal potential from the intersection of the linear current–voltage curve with the x-axis.

We performed whole-cell patch clamp recordings from CA1 pyramidal neurons and GAD65-GFP interneurons in the *stratum Radiatum* (sRad) using borosilicate glass pipettes with resistances of 3-6 MΩ. For recordings of spontaneous inhibitory postsynaptic currents (sIPSCs), miniature inhibitory currents (mIPSCs) and evoked IPSCs (eIPSCs), pipettes were filled with a KCl-based internal solution (containing in mM: 70 KGluconate, 70 KCl, 0.5 ethyleneglycol-bis(β-aminoethyl)-N,N,N’,N’-tetraacetic Acid (EGTA), 4 Na_2_phosphocreatine, 4 MgATP, 0.4 NaGTP and 10 HEPES; pH adjusted to 7.2 with KOH) and ACSF was supplemented with 20 μM DNQX (Tocris) and 50 μM DL-APV (Tocris). For recordings of miniature IPSCs (mIPSCs), 1 μM TTX (Abcam) was also added to the ACSF. For recordings of spontaneous excitatory postsynaptic currents (sEPSCs), pipettes were filled with a K-gluconate-based internal solution (containing in mM: 140 K-gluconate, 4 KCl, 0.5 EGTA, 10 HEPES, 4 MgATP, 0.4 NaGTP, 4 Na_2_phosphocreatine, 30 Alexa 568 (Thermo Fisher Scientific); pH adjusted to 7.2 with KOH) and ACSF was supplemented with 6 μM gabazine (HelloBio). Membrane potential was clamped at -65 mV. sIPSCs and mIPSCs were recorded for 6 minutes, sEPSCs for 10 minutes. For recording of eIPSCs a glass electrode filled with ACSF and containing a stimulus electrode was positioned either in sRad or *stratum Pyramidale* (sPyr) of CA1 (respectively ∼ 250–300 μm or ∼50 μm from the recording electrode). Stimulus intensity was set at half maximum response. eIPSCs were recorded at least 30 times every 30s with 100 ms interval. Paired pulse ratios were calculated as the mean amplitude of all second responses divided by the mean amplitude of all first responses (Kim and Alger, 2001). The coefficient of variation was calculated as the standard deviation divided by the mean amplitude of the first responses (n=15-60). After recording of sEPSCs, we switched to current clamp to assess excitability. Current was injected at resting membrane potential (I=0) in 10 pA increments with a maximum of 200 pA to assess firing properties. The threshold potential was determined as the accompanying membrane potential at the start of the action potential. Cells were discarded if series resistance was above 35 MΩ or if the resting membrane potential exceeded -50 mV for pyramidal neurons and -40 mV for GAD65-GFP interneurons or if the series resistance after the recording deviated more than 30% from its original value.

Cell-attached recordings were performed using borosilicate glass pipettes of 3-5 MΩ, filled with a 150 mM NaCl-based solution. Modified ACSF (Maffei et al., 2004) was used to increase the baseline firing frequency (containing in mM: 126 NaCl, 3,5 KCl, 1.0 CaCl_2_, 0.5 MgCl_2_, 26 NaHCO_3_, 1.25 Na_2_H_2_PO_4_, 20 glucose). CA1 pyramidal neurons were sealed (≥1 GΩ) and voltage clamped at a holding current of 0 pA, to avoid affecting the firing activity of the cell (Perkins, 2006). Firing was recorded for 6 minutes before and after wash in of muscimol (50 μM, Tocris).

All data were blinded before analysis. Network discharges were marked and removed from sEPSC recordings before event analysis. Events were selected using a template in Clampfit10 software. Further analysis was performed using custom-written MATLAB scripts. Rise time of sIPSCs was determined as the time between 10% and 90% of the peak value. The distribution of the rise times recorded in control conditions (generated from 100 randomly selected IPSCs per cell) was fitted with two Gaussians and their crossing point determined the separation between fast and slow IPSCs (Ruiter et al., 2020). We verified that the double Gaussian fit for the rise time distribution in VU conditions gave a similar separation value (control: 0.85 ms; VU: 0.95 ms) and our conclusions did not change by taking the VU separation value. The decay tau was fitted with a single exponential function. Only events with a goodness of fit R^2^ ≥ 0.75 were included.

### Two-photon SuperClomeleon imaging and analysis

We performed two-photon chloride imaging in CA1 pyramidal cells using the SuperClomeleon sensor (Grimley et al., 2013) in cultured slices from SClm mice (described above). To target GABAergic interneurons, we used an AAV approach in slices from VGAT-Cre mice. An adeno associated virus (AAV) 2/5 for Cre-dependent SClm expression was prepared from a SCLM-DIO construct (kind gift from Thomas Kuner (Boffi et al., 2018)) using packaging plasmid pAAV2/5 (addgene_104964) and helper plasmid pAdDeltaF6 (addgene_112867). The virus was added on top of the CA1 region of organotypic hippocampal cultures from VGAT-Cre mice at DIV2 using a microinjector (Eppendorf, FemtoJet) aided by a stereoscopic microscope (Leica, M80). This resulted in widespread, but sparse SClm expression in GABAergic neurons.

At the recording day, slices were transferred to a recording chamber and continuously perfused with carbonated (95% O_2_, 5% CO_2_) ACSF supplemented with 1 mM Trolox, at a rate of approximately 1 ml/min. Bath temperature was monitored and maintained at 30-32 °C. Two-photon imaging of CA1 pyramidal neurons or VGAT-positive interneurons in the CA1 area of the hippocampus was performed using a customized two-photon laser scanning microscope (Femto3D-RC, Femtonics, Budapest, Hungary) with a Ti-Sapphire femtosecond pulsed laser (MaiTai HP, Spectra-Physics) and a 60x water immersion objective (Nikon NIR Apochromat; NA 1.0) The CFP donor was excited at 840 nm. The emission light was split using a dichroic beam splitter at 505 nm and detected using two GaAsP photomultiplier tubes. We collected fluorescence emission of Cerulean/CFP (485 ± 15 nm) and Topaz/YFP (535 ± 15 nm) in parallel. Of each slice from SClm mice, 2-5 image stacks were acquired in different fields of view. Of each VGAT-Cre slice 2-5 image stacks were acquired in different fields of view in both sPyr and sRad).The resolution was 8.1 pixels/μm (1024×1024 pixels, 126×126 μm) with 1 μm z-steps.

Image analysis was performed using ImageJ software, as described before (Herstel et al., 2022). We manually determined regions of interest (ROIs) around individual neuron somata. To select a representative cell population in SClm slices, in each image z-stack we selected four z-planes at comparable depths in which three pyramidal cells were identified that varied in CFP brightness (bright, middle and dark). In VGAT-Cre slices with viral SClm expression, all visible neurons were analyzed (range: 1 to 9). We subtracted the mean fluorescence intensity of the background in the same image plane before calculating the fluorescence ratio of CFP and YFP. We limited our analysis to cells which were located within 450 pixels from the center of the image, as FRET ratios showed slight aberrations at the edge of our images. We excluded cells with a FRET ratio < 0.5 or > 1.6 to avoid unhealthy cells. FRET-colored images (as shown in Fig. 2D and 9A) were made in ImageJ. We first subtracted the average background and Gaussian filtered the CFP and YFP image z-stacks separately. Next, the acceptor (YFP) image was divided by the donor (CFP) image to get the ratiometric image. A mask was created by manually drawing ROIs for each soma in the image. An average projection of the ratiometric image was made in a specific z range and multiplied by the mask. Finally, the masked ratiometric image was combined with the grayscale image. Please note that these images were made for illustration purposes only, analysis was done on the raw data.

**Fig. 1.**
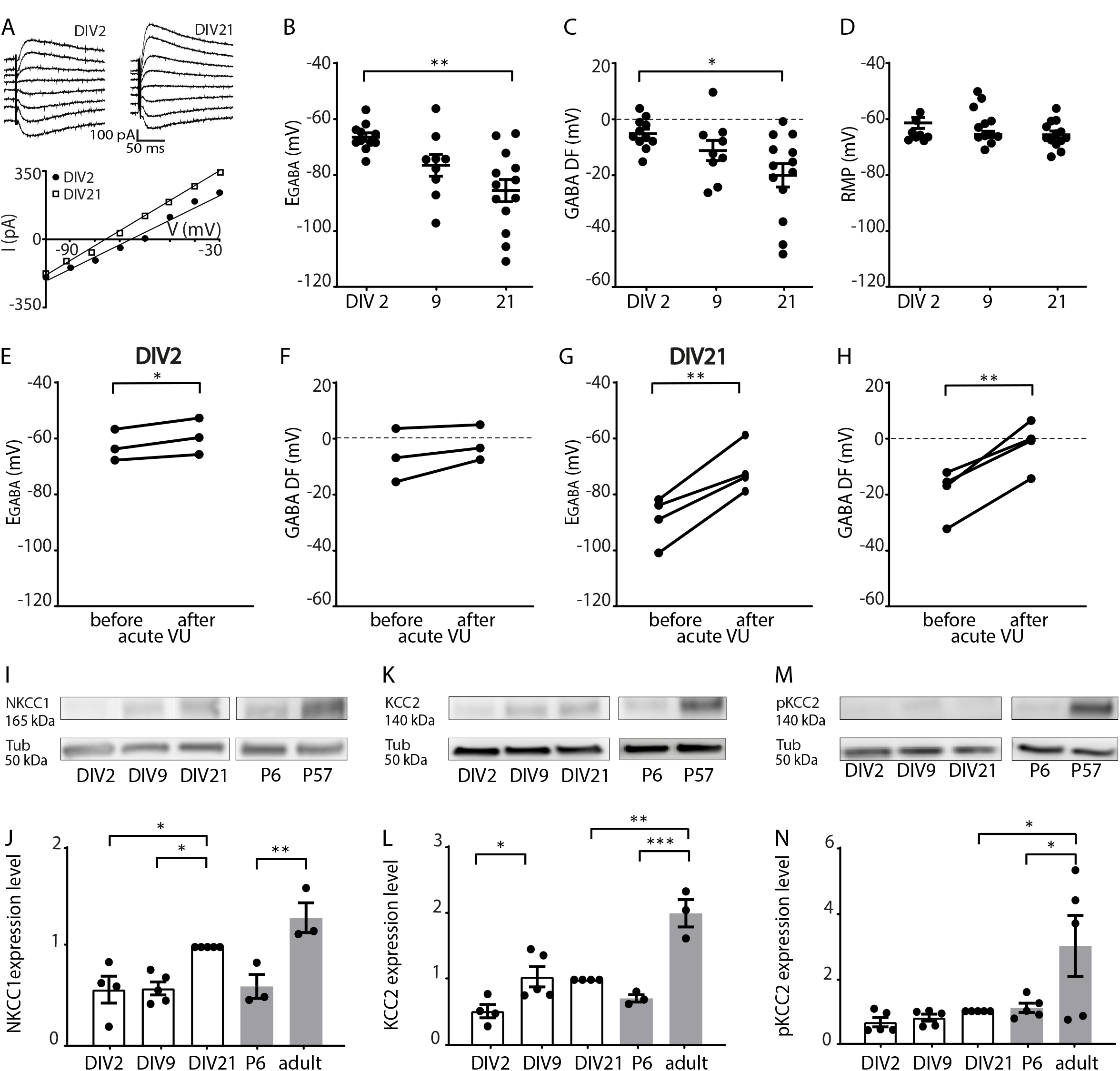
GABA shift is ongoing in organotypic hippocampal slice cultures. A) Perforated patch clamp recordings in CA1 pyramidal cells in hippocampal slice cultures at DIV2 and DIV21. Responses to muscimol application were recorded at holding potentials from -100 to -30 mV with 10 mV increments and GABA reversal potential was determined from the intersection of the linear current-voltage curve with the x-axis. B) The GABA reversal potential (E_GABA_) recorded in cultured slices at DIV2, DIV9 and DIV21 (KW, p=0.019; DMC: DIV2 versus DIV9, p=0.13; DIV9 versus DIV 21, p=0.65; DIV2 versus DIV21, p=0.001). C) GABA Driving Force (GABA DF) during *in vitro* slice development (KW, p=0.017; DMC: DIV2 versus DIV9, p=0.35; DIV9 versus DIV21, p=0.85; DIV2 versus DIV21, p=0.013). D) The resting membrane potential (RMP) during *in vitro* slice development (KW, p=0.23). Data in B-D from 9-13 cells from 8-11 slices and 5-10 mice per group. E,F) E_GABA_ (PT, p=0.038) and GABA DF (PT, p=0.18) values recorded in CA1 pyramidal cells at DIV2 before and after acute application of VU. Data from 3 cells, 3 slices and 3 mice per group. G,H) E_GABA_ (PT, p=0.009) and GABA DF (PT, p=0.006) values recorded in CA1 pyramidal cells at DIV21 before and after acute application of VU. Data from 4 cells, 4 slices and 4 mice per group. I) Western blot for NKCC1 protein content, measured in hippocampal slices cultures at DIV2, DIV9 and DIV21, and in acute slices from P6 and adult (P57) mice. Tubulin (Tub) was used as loading control. J) Summary of data for NKCC1 expression levels. Protein levels were normalized to DIV21 values. (1W ANOVA, p=0.0003; SMC: DIV2 versus DIV9, p>0.99; DIV9 versus DIV21, p=0.019; DIV2 versus DIV21, p=0.023; DIV21 versus P57, p=0.26; P6 versus P57, p=0.003). K) Western blot for KCC2 protein content, measured in hippocampal slices cultures at DIV2, DIV9 and DIV21, and in acute slices from P6 and adult (P57) mice. Tubulin (Tub) was used as loading control. L) Summary of data for KCC2 expression levels. Protein levels were normalized DIV21 values. (1W ANOVA, p<0.0001; SMC: DIV2 versus DIV9, p=0.045; DIV9 versus DIV21, p>0.99; DIV2 versus DIV21, p=0.10; DIV21 versus P57, p=0.0007; P6 versus P57, p=0.0001). M) Western blot for S940-pKCC2 (pKCC2) protein content, measured in hippocampal slices cultures at DIV2, DIV9 and DIV21, and in acute slices from P6 and adult (P57) mice. Tubulin (Tub) was used as loading control. N) Summary of data for s940-pKCC2 (pKCC2) expression levels. Protein levels were normalized to DIV21 values. (1W ANOVA, p=0.006; SMC: DIV2 versus DIV9, p>0.99; DIV9 versus DIV21, p>0.99; DIV2 versus DIV21, p=0.99; DIV21 versus P57, p=0.020; P6 versus P57, p=0.033). Data in I-N from 3-5 experiments and 2-5 mice per group.

**Fig. 2.**
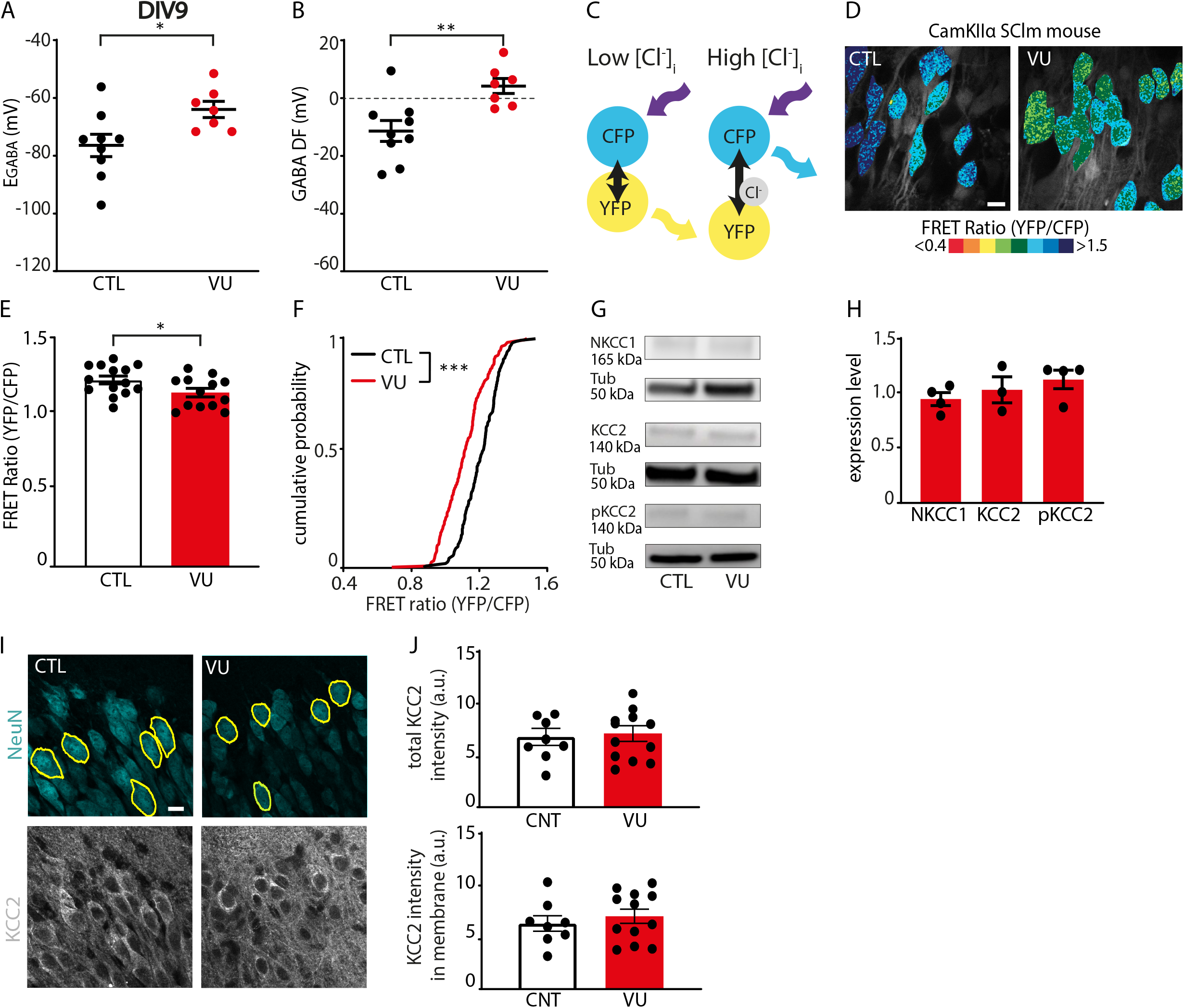
VU depolarizes GABA DF and increases intracellular chloride levels at DIV9. A,B) GABA reversal potential (E_GABA_) (MW, p=0.016) and GABA Driving Force (GABA DF) (MW, p=0.005) recorded in CA1 pyramidal cells at DIV9 in control (CTL) and VU-treated slices. Data from 7-9 cells, 6-8 slices and 3-5 animals per group. C) Illustration of Fluorescence Resonance Energy Transfer (FRET) from CFP donor to YFP acceptor of the SClm sensor. FRET ratios (YFP/CFP fluorescence) decrease with increasing chloride concentrations. D) Two-photon images of SClm FRET ratios in CA1 pyramidal neurons in CTL and VU-treated cultures at DIV9. Individual cells are color-coded to their FRET ratios. Scale bar: 10 μm. E) Average SClm FRET ratios in CTL and VU-treated cultures (UT, p=0.041). Data from 16-24 cells per slice, 13-14 slices and 6-8 mice per group. F) Cumulative distribution of FRET ratios in individual cells in CTL and VU-treated cultures. (KS, p<0.0001). Data from 15 cells per slice, 13-14 slices and 6-8 mice per group. G) Western blot of NKCC1, KCC2 and S940-pKCC2 (pKCC2) expression in CTL and VU-treated cultures. Tubulin (Tub) was used as loading control. H) Summary of data for NKCC1, KCC2 and for s940-pKCC2 (pKCC2) expression levels at DIV9. Values in VU-treated cultures were normalized to the protein level in CTL cultures (NKCC1: MW, p>0.99; KCC2: MW, p=0.70; pKCC2: MW, p=0.31). I) Confocal images of NeuN and KCC2 staining in CTL and VU-treated cultures with ROIs in yellow. Scale bar: 10 μm. J) Total KCC2 levels (UT, p=0.78) and KCC2 levels in membrane (UT, p=0.51) in CTL and VU-treated cultures at DIV9. Each datapoint represents the mean KCC2 intensity of the 5-15 ROIs in one image. Data from 8-12 images, 4-6 slices and 3 mice per group.

### Protein extraction and Western Blot analysis

Organotypic hippocampal cultures were washed in cold PBS and subsequently lysed in cold protein extraction buffer containing: 150 mM NaCl, 10 mM EDTA 10 mM HEPES, 1% Triton-X100 and 1x protease and 1x phosphatase inhibitors cocktails (Complete Mini EDTA-free and phosSTOP, Roche). Lysates were cleared of debris by centrifugation (14,000 rpm, 1 min, 4°C) and measured for protein concentration before storage at −20°C until use. Lysates were denatured by adding loading buffer and heating to 95°C for 5 min. For each sample an equal mass of proteins was resolved on 4–15 % polyacrylamide gel (Bio-Rad). The proteins were then transferred (300 mA, 3h) onto ethanol-activated Immobilon-P PVDF membrane (Millipore) before blocking with 3% Bovine Serum Albumin in Tris-buffered saline-Tween (TBST, 20 mM Tris, 150 mM NaCl, 0.1% Tween-20) for 1 h. Primary antibodies used in this study were: mouse anti-NKCC1 (T4, Developmental Studies Hybridoma Bank, 1:1000), rabbit anti-KCC2 (07-432, Merck, 1:1000) and s940-pKCC2 (p1551-940, LuBioSciences, 1:1000), mouse anti-tubulin (T-5168, Sigma, 1:10000). Primary antibodies were diluted in blocking buffer and incubated with the blots overnight at 4 °C under gentle rotation. The membrane was washed 3 times 15 minutes with TBST before a 1h incubation of horseradish peroxidase (HRP)-conjugated antibodies (P0447 goat anti-mouse IgG HRP, Dako, 1:2500 or P0399 swine anti-rabbit IgG HRP, Dako, 1:2500), and washed 3 times 15 minutes in TBST again before chemiluminescence detection. For chemiluminescence detection, blots were incubated with Enhanced luminol-based Chemiluminescence substrate (Promega), and the exposure was captured using the ImageQuant 800 system (Amersham). Images were analyzed in ImageJ, by drawing rectangular boxes around each band and measuring average intensities. Protein levels were normalized to the Tubulin loading controls.

### Immunohistochemistry, confocal microscopy and analysis

Organotypic hippocampal cultures were fixed 4 % paraformaldehyde solution in PBS for 30 minutes at room temperature. After washing slices in phosphate buffered saline (PBS), they were permeabilized for 15 minutes in 0.5 % Triton X-100 in PBS for 15 minutes, followed by 1 hour in a blocking solution consisting of 10 % normal goat serum and 0.2 % Triton X-100 in PBS. Slices were incubated in primary antibody solution at 4 °C overnight. The following primary antibodies were used: rabbit polyclonal anti-KCC2 (07-432, Merck, 1:1000), rabbit anti-VGAT (131003, Synaptic Systems, 1:1000), guinea pig anti-VGLUT (AB5905, Merck, 1:1000), mouse anti-NeuN (MAB377, Millipore, 1:500), guinea pig anti-NeuN Merck, 1:1000).

Slices were washed in PBS and incubated in secondary antibody solution for four hours at room temperature. Secondary antibodies used were: goat anti-mouse Alexa Fluor 467 (Life Technologies, A21236, 1:500), goat anti-rabbit Alexa Fluor 405 (Life Technologies, A31556, 1:500), goat anti-guinea pig Alexa Fluor 488 (A11073, Life Technologies, 1:1000) and Goat anti-guineapig Alexa Fluor 568 (A11075, Life Technologies, 1:500) Goat anti-rabbit Alexa fluor 647 (A21245, Life Technologies, 1:500). After another PBS wash, slices were mounted in Vectashield mounting medium (Vector labs).

Confocal images were taken on a Zeiss LSM-700 confocal laser scanning microscopy system with a Plan-Apochromat 40x 1.3 NA immersion objective for KCC2 staining and 63x 1.4 NA oil immersion objective for spines and VGLUT/VGAT staining. VGLUT/VGAT staining was imaged in the CA1 sPyr and sRad of each slice, KCC2 staining was imaged in two fields of view in the CA1 sPyr of each slice and spines were imaged on the second dendritic branch of the apical dendrite of each Alexa-568 filled neuron. Image stacks (1024×1024 pixels) were acquired at 6.55 pixels/μm for KCC2 staining, 20.15 pixels/μm for spines and 10.08 pixels/μm for VGLUT/VGAT staining. Step size in z was 0.5 μm for spines and KCC2 staining and 0.4 μm for VGLUT/VGAT staining.

Confocal images were blinded before analysis. The number of dendritic spines on the second dendritic branch of the apical dendrite were counted using the multi-point tool in ImageJ. As we noticed that spine density was low near the base of dendrites, the first 10 μm of the dendrites were excluded from analysis. For analysis of VGLUT and VGAT puncta a custom-made macro was used (Ruiter et al., 2020). Briefly, three z-planes were averaged and background was subtracted with rolling ball radius of 10 pixels. Puncta were then identified using watershed segmentation. For the analysis of KCC2 levels, ROIs were drawn around 5 NeuN positive cell bodies, and this was repeated in 1-3 z-planes (depending on the depth of KCC2 signal) at comparable depths, to measure KCC2 fluorescence in 5-15 neurons per image stack. We determined average fluorescent intensities in the ROI as well as within the ROIs after scaling it with 10 pixels (∼1 um) in width to assess both the total and membrane fraction of KCC2.

### Experimental design and statistical analyses

Statistical analysis was performed in Prism (Graphpad). Normality was tested using the D’Agostino & Pearson test. For the comparison of two groups we either used an unpaired Student’s t test (UT; parametric), a Mann-Whitney test (MW; non-parametric), a paired Student’s t test (PT; parametric) or a Wilcoxon signed-rank test (WSR; non-parametric). For comparison of two cumulative distributions we performed a Kolmogorov Smirnov test (KS; nonparametric). For comparison of multiple groups, a one way ANOVA (1W ANOVA; parametric) was used, followed by a by a Sidak’s Multiple Comparisons posthoc test (SMC) or a Kruskal Wallis test (KW; nonparametric) was used, followed by a Dunn’s Multiple Comparison posthoc test (DMC). For comparison of two variables in multiple groups a two way ANOVA (2W ANOVA; parametric) was performed, followed by a Sidak’s Multiple Comparisons posthoc test (SMC). Data are presented as mean ± standard error of the mean. Significance is reported as *p<0.05; **p≤0.01; ***p≤0.001.

## Results

### The GABA shift takes place in organotypic hippocampal cultures

We assessed the development of neuronal chloride levels in organotypic hippocampal cultures made from P6-7 mouse pups using gramicidin perforated patch recordings of CA1 pyramidal neurons. We recorded GABAergic currents in response to brief applications of the GABA_A_ receptor agonist muscimol and determined the GABA reversal potentials at DIV2, DIV9 and DIV21 (Fig. 1A). The GABA reversal potential (E_GABA_) and GABAergic driving force (GABA DF) decreased gradually over this period (Fig. 1B, C), indicating that intracellular chloride homeostasis was still maturing in our slice cultures (Salmon et al., 2020; Herstel et al., 2022). The resting membrane potential (RMP) remained stable (Fig. 1D).

Intracellular chloride levels are determined by the relative expression level and activity of the chloride importer NKCC1 and the chloride exporter KCC2 (Rivera et al., 1999; Gulyás et al., 2001; Yamada et al., 2004; Banke and McBain, 2006; Otsu et al., 2020). We therefore determined the contribution of KCC2 to the GABA reversal potential. Application of KCC2 blocker VU0463271 (VU) in slices at DIV2 resulted only in a small depolarizing shift of E_GABA_ (Fig. 1E), and did not affect GABA driving force (Fig. 1F), while in DIV21 slices VU application triggered an acute elevation of E_GABA_ and GABA driving force (Fig. 1G, H). This suggests that KCC2 function increased with development *in vitro*, as expected (Salmon et al., 2020). We assessed the expression levels of NKCC1 and KCC2 in our slice cultures using Western blots. As the function of KCC2 is enhanced during development by phosphorylation at serine 940 (S940) (Lee et al., 2007; Mòdol et al., 2014), we also included a phospho-specific antibody to detect phosphorylated KCC2 (pKCC2) levels. The expression of NKCC1 and KCC2 increased in our organotypic cultures between DIV2 and DIV21 (Fig. 1I-L). For comparison with the developmental trajectory *in vivo*, we determined NKCC1, KCC2 and pKCC2 levels in hippocampal tissue from P6 and adult mice. NKCC1, KCC2 and pKCC2 expression increased from P6 to adulthood (Fig. 1I-N). We noted that KCC2 and pKCC2 expression in DIV21 slice cultures did not reach adult levels (Fig. 1K-N). Together, these data show that a clear GABA shift occurred *in vitro*, which is consistent with a previous report (Salmon et al., 2020) and our own previous analysis of chloride maturation in organotypic hippocampal cultures (Herstel et al., 2022).

### VU treatment depolarizes GABA driving force and elevates intracellular chloride levels at DIV9

Having established that the GABA shift occurs during the first week *in vitro*, we set out to delay the shift with one week to mimic the delayed chloride development in NDD mouse models (He et al., 2014; Tyzio et al., 2014; Deidda et al., 2015b; Banerjee et al., 2016; Corradini et al., 2017; Fernandez et al., 2018; Roux et al., 2018; Lozovaya et al., 2019; Bertoni et al., 2021). We treated slices for one week (from DIV2 to DIV8) with VU0463271 (VU), a specific blocker of the chloride exporter KCC2 (Delpire et al., 2012), or with DMSO control. One week treatment with VU from resulted in a significant elevation of E_GABA_ in CA1 pyramidal neurons immediately after the VU treatment at DIV9 (Fig. 2A). The driving force for GABA signaling (GABA DF) also shifted to positive values (Fig. 2B), which were comparable to control slices at DIV2 (Fig. 1C). This indicates that GABA signaling remained depolarizing in VU-treated slices at DIV9. We assessed intracellular chloride levels in the CA1 pyramidal cell population using the chloride sensor SuperClomeleon (SClm) (Grimley et al., 2013; Boffi et al., 2018; Rahmati et al., 2021). The SClm sensor consists of two fluorescent proteins, Cerulean (CFP mutant) and Topaz (YFP mutant), joined by a flexible linker. Depending on the binding of chloride, Fluorescence Resonance Energy Transfer (FRET) occurs from the CFP donor to the YFP acceptor ((Grimley et al., 2013); Fig. 2C). We measured SClm FRET ratios (fluorescence intensity of YFP/CFP) using two-photon microscopy. Measured FRET ratios are independent of expression level and tissue depth (Herstel et al., 2022). We observed a prominent decrease in SClm FRET values in CA1 pyramidal neurons (Fig. 2D,E), reflecting an increase in chloride levels after VU treatment. When we analyzed the distribution of individual values we observed a clear increase in the fraction of cells with low FRET ratios (Fig. 2F). Together, these data show that VU treatment between DIV2 and DIV8 resulted in an elevation of intracellular chloride levels and depolarizing GABA signaling at DIV9.

Next, we assessed whether the expression levels of the chloride cotransporters NKCC1 and KCC2 were altered after the VU treatment. We did not find any significant difference in the levels of NKCC1, KCC2 or S940-pKCC2 between VU-treated and control slices at DIV9 (Fig. 2G,H). As Western blot only allows for assessment of total KCC2 levels and cannot distinguish between cytosolic KCC2 and KCC2 in the plasma membrane, we also examined KCC2 levels in treated and control slices using immunohistochemistry. We used NeuN to identify individual cell bodies and we estimated KCC2 levels in these cells. VU treatment did not significantly affect total or membrane KCC2 levels (Fig. 2I,J). Together, this indicates that blocking KCC2 function between DIV2 and DIV8 delayed the GABA shift without changing the expression or surface levels of KCC2.

### Synaptic inputs and firing properties of CA1 pyramidal cells are not affected at DIV9 after VU treatment

Previous experimental studies in which the GABA shift was accelerated showed that depolarizing GABA facilitates excitatory synapse formation (Leinekugel et al., 1997; Akerman and Cline, 2006; Wang and Kriegstein, 2008, 2011; Chancey et al., 2013; van Rheede et al., 2015; Oh et al., 2016). This would predict that delaying the GABA shift, e.g. prolonging the period when GABA signaling is depolarizing, would result in an excess synapse growth.

To assess functional synapses, we recorded sEPSCs and sIPSCs in CA1 pyramidal cells using whole-cell patch clamp recordings (Fig. 3A,B). We found that sEPSC frequency, amplitudes, and kinetics were not different in CA1 pyramidal cells in VU-treated and control slices at DIV9 (Fig. 3C-F). There were also no changes in sIPSCs (Fig. 3G-J). In addition, we recorded action potential (AP) firing rates during current injections of increasing amplitude (Fig. 3K). Firing rates, resting membrane potential and action potential threshold were comparable in control and VU-treated slices (Fig. 3L-N).

**Fig. 3.**
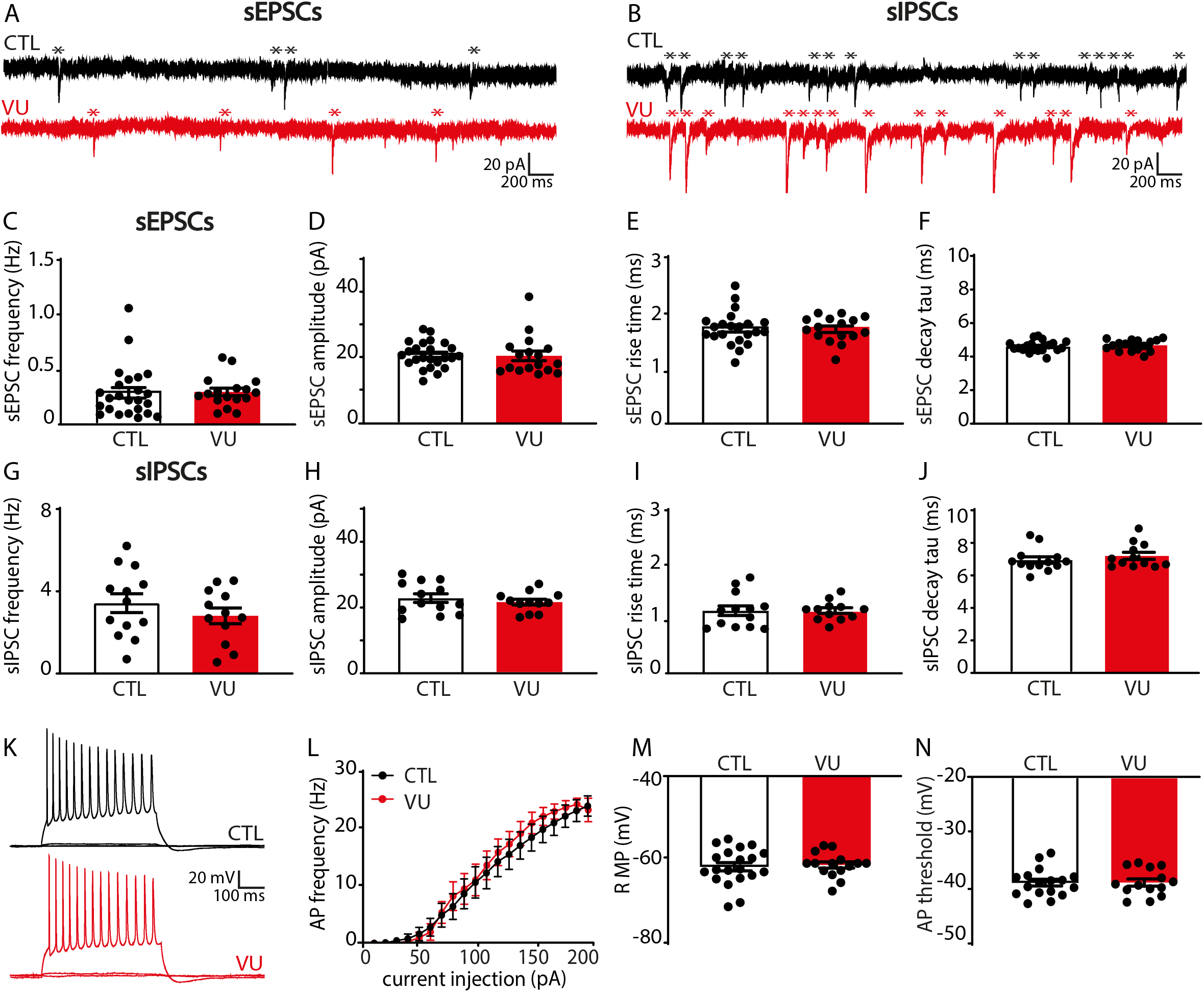
VU treatment does not change excitatory or inhibitory transmission and firing properties at DIV9. A) sEPSC recordings from CA1 pyramidal cells in a control (CTL) and VU-treated culture at DIV9. sEPSCs are indicated with asterisks. B) sIPSC recordings in CTL and VU-treated cultures at DIV9. sIPSCs are indicated with asterisks. C-F) sEPSC frequency (MW, p=0.36), amplitude (MW, p=0.45), risetime (UT, p=0.87) and decay tau (UT, p=0.53) in CTL and VU-treated cultures at DIV9. Data from 16-24 cells, 11-15 slices and 10-12 mice per group. G-J) sIPSC frequency (UT, p=0.32), amplitude (UT, p=0.46), risetime (UT, p=0.99) and decay tau (UT, p=0.20) in in CTL and VU-treated cultures at DIV9. Data from 12-13 cells, 4-7 slices and 4-6 mice per group. K) Example recordings of action potentials during current injections in CA1 pyramidal neurons in CTL and VU-treated cultures at DIV9. L) Action potential firing rates in CTL and VU-treated cultures with increasing current injections at DIV9 (2W ANOVA, current injection: p<0.001, treatment: p=0.24). Data from 14-17 cells, 10-11 slices and 9-10 mice per group. M,N) Resting membrane potential (RMP) (UT, p=0.75) and action potential (AP) threshold (UT, p=0.98) in CTL and VU-treated cultures at DIV9. Data from 14-20 cells, 10-11 slices and 9-10 mice per group.

Additionally, we assessed synaptic density by performing immunohistochemistry on VU-treated and control slices. Consistent with our electrophysiological data, the density and size of VGLUT and VGAT puncta in the *stratum Pyramidale* (sPyr) and *stratum Radiatum* (sRad) of the CA1 area were similar in VU-treated and control slices at DIV9 (Fig. 4A-J). Furthermore, we filled pyramidal neurons with Alexa-568 via a patch pipette to assess dendritic spines at which excitatory synapses are located (Sheng and Hoogenraad, 2007). Spine density of CA1 pyramidal neurons was similar in control and VU-treated slices (Fig. 4K,L).

**Fig. 4.**
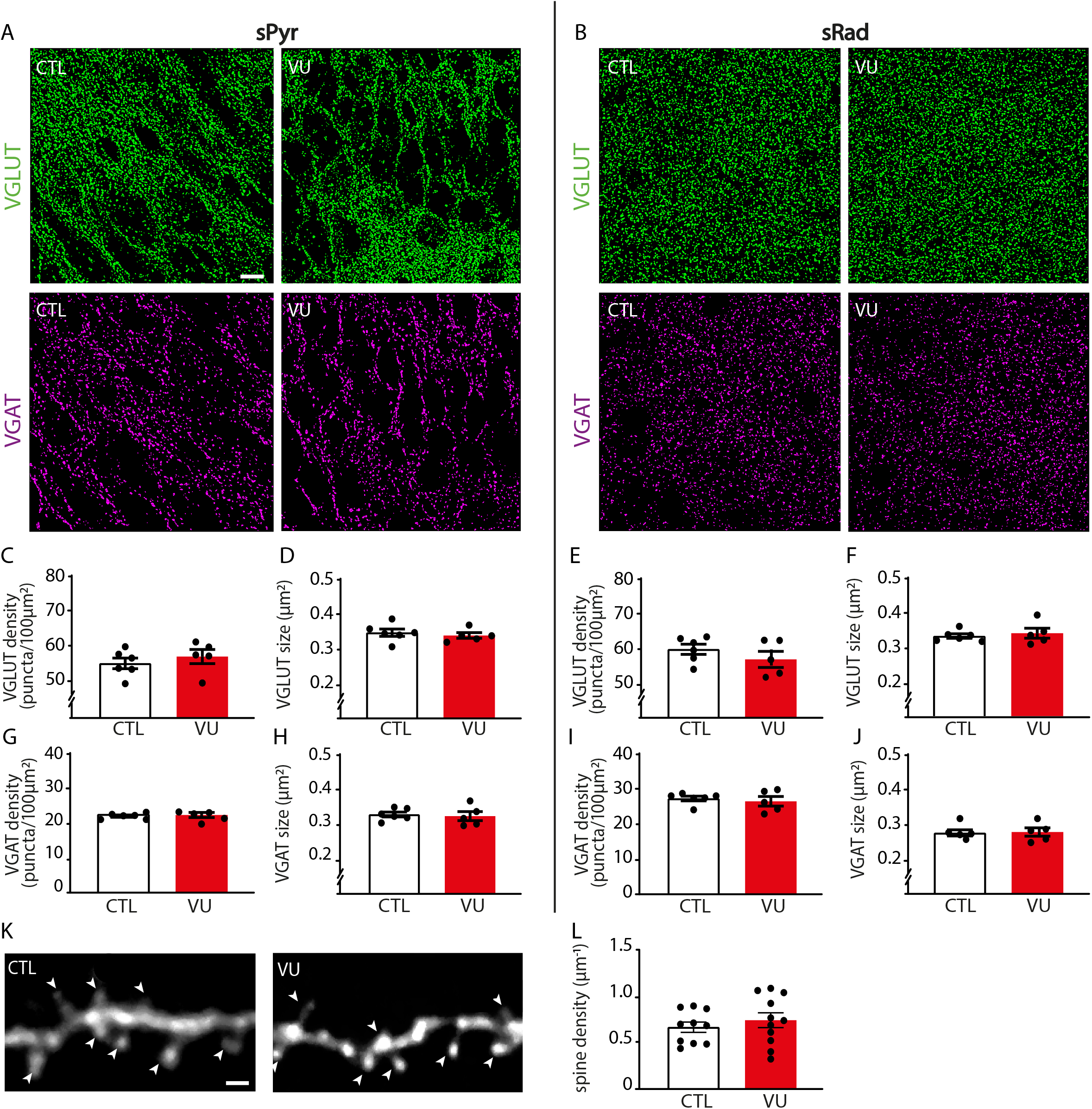
VU treatment does not change excitatory or inhibitory synapses at DIV9. A,B) Thresholded images of VGLUT and VGAT puncta in the CA1 sPyr and sRad of control (CTL) and VU-treated cultures at DIV9. Scale bar: 10 μm. C,D) Density (MW, p=0.43) and size (MW, p=0.33) of VGLUT puncta in the sPyr in CTL and VU-treated cultures at DIV9. E,F) Density (MW, p=0.27) and size (MW, p=0.83) of VGLUT puncta in the sRad. G,H) Density (MW, p=0.96) and size (MW, p=0.54) of VGAT puncta in the sPyr. I,J) Density (MW, p=0.89) and size (MW, p=0.84) of VGAT puncta in the sRad. Data in C-J from 5-6 slices and 4-5 mice per group. K) Example images of the apical dendrite of CA1 pyramidal neurons in CTL and VU-treated cultures at DIV9. Spines are indicated with arrowheads. Scale bar: 1 μm. L) The average density of dendritic spines (UT, p=0.82) in CTL and VU-treated cultures at DIV9. Data from 10-11 cells, 6-8 slices and 6 mice per group.

Together, these data show that VU treatment from DIV2 to DIV8 induced a clear depolarizing shift in GABA signaling, but this did not affect excitatory and inhibitory synapses and firing properties in CA1 pyramidal cells at DIV9.

### Depolarizing GABA signaling in VU-treated slices acts inhibitory

As eluded to above, it is important to note that depolarizing GABA signaling does not necessarily mean that GABA acts inhibitory. Depolarizing GABA can have an inhibitory action and limit neuronal activity when the GABA-mediated depolarization is subthreshold and shunts excitatory inputs (Staley and Mody, 1992; Kirmse et al., 2015; Murata and Colonnese, 2020; Salmon et al., 2020). We noticed that during the sEPSC recordings, when GABA_A_-mediated inhibitory currents were blocked with gabazine, network discharges often occurred. These discharges probably reflect periods of synchronous firing of pyramidal cells that can occur in the absence of fast GABA_A_-mediated inhibition, when only the slower GABA_B_-mediated inhibition is left to counteract excitatory synaptic transmission (Scanziani et al., 1994; Menendez De La Prida et al., 2006). The frequency and duration of these network discharges were not different in VU-treated and control slices at DIV9 (Fig. 5A-C). We directly assessed if depolarizing GABA was already inhibitory in our VU-treated slices. To determine the role of GABA in regulating action potential firing, we washed in GABA_A_ receptor agonist muscimol while recording spontaneous firing in cell-attached configuration. As spontaneous activity of hippocampal neurons is very low in standard artificial cerebral spinal fluid (ACSF), we used a modified ACSF, which elevates spontaneous firing (Maffei et al., 2004). Muscimol inhibited firing in control as well as in VU-treated slices at DIV8 (Fig. 5D-F). This indicates that GABA_A_ receptor-mediated signaling is inhibitory in both control and VU-treated slices at DIV8, despite being hyperpolarizing in control and depolarizing in VU-treated slices (Fig. 2B). This suggests that GABA signaling mostly acts inhibitory in hippocampal slice cultures from P6-7 mice, which is in line with previous reports in P6-9 acute hippocampal slices (Khalilov et al., 1997; Valeeva et al., 2016), cultured hippocampal slices at P5 DIV3-5 (Salmon et al., 2020) and *in vivo* recordings in the hippocampus after P3-7 (Valeeva et al., 2010; Murata and Colonnese, 2020). Together, our data suggest that depolarizing GABA signaling after the first postnatal week acts mostly inhibitory and does not contribute to synapse formation in the hippocampus.

**Fig. 5.**
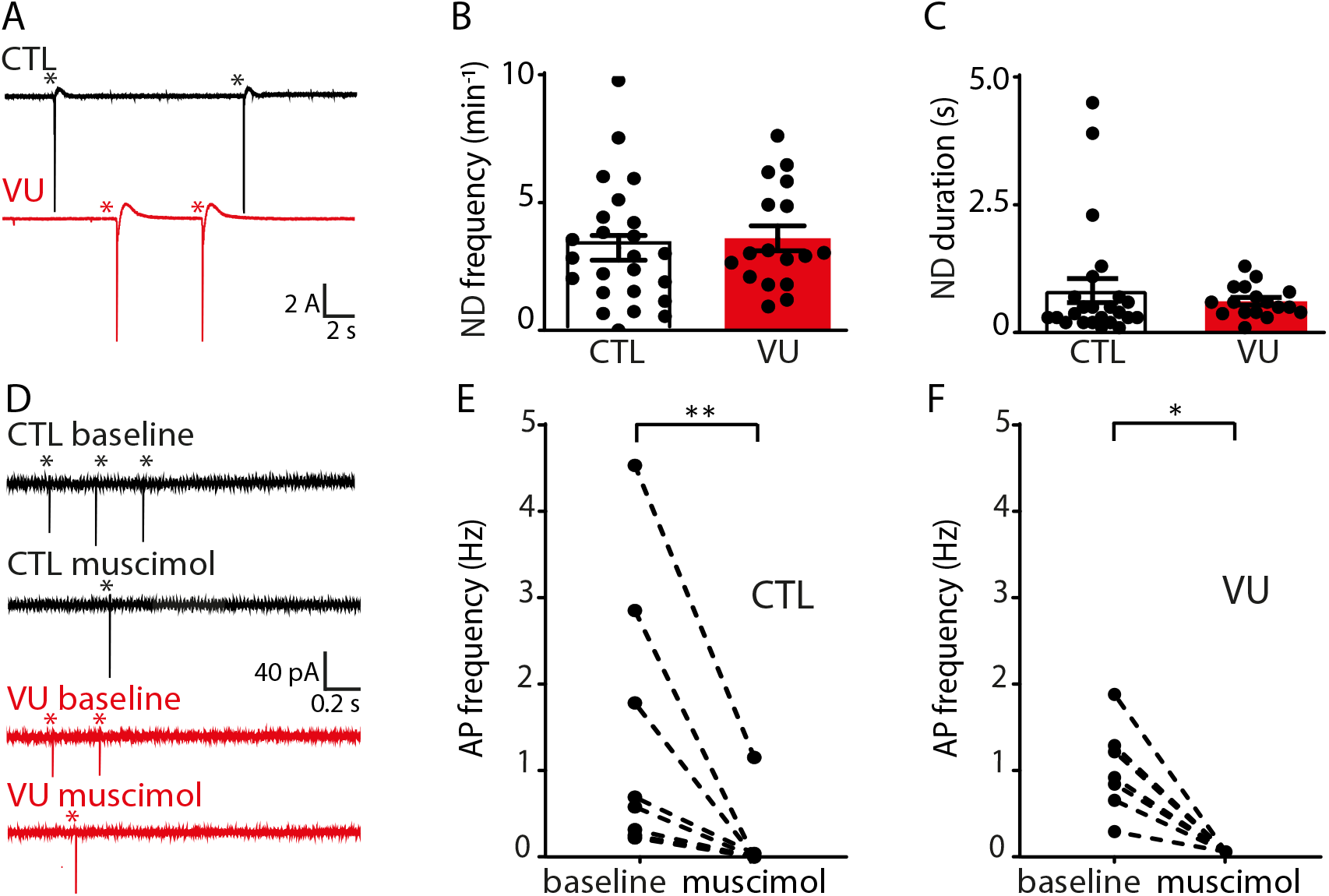
GABA is inhibitory in control and VU-treated slice cultures at DIV8 and DIV9. A) Whole-cell voltage clamp recordings from CA1 pyramidal cells in the presence of gabazine in control (CTL) and VU-treated cultures at DIV9. Network discharges are indicated with asterisks. B,C) Network discharge (ND) frequency (MW, p=0.61) and duration (MW, p=0.20) in CTL and VU-treated cultures at DIV9. Data from 17-24 cells, 11-15 slices and 10-12 mice per group. D) Cell-attached recordings from CA1 pyramidal cells showing action potential (AP) firing in modified ACSF in CTL and VU-treated cultures at DIV8 before (baseline) and after muscimol wash in. APs are indicated with asterisks. E, F) Average AP frequency before and after muscimol washin in CTL and VU-treated cultures at DIV8. Muscimol decreases firing rates in CTL (WSR, p=0.008) and VU-treated (WSR, p=0.016) cultures at DIV8. Data from 7-8 cells, 7-8 slices and 3-4 mice per group. A MW was performed to compare baseline firing rates in CTL and VU-treated cultures (MW, p=0.79).

### Indirect effects on inhibitory synaptic inputs in VU-treated slices at DIV21

Previous studies demonstrated that transient alterations in GABA function during early postnatal development can have long-lasting consequences for network function and ultimately behavior (He et al., 2018; Bertoni et al., 2021; Matsushima et al., 2022). We therefore assessed possible long-lasting consequences of the VU treatment on the CA1 network at DIV21. After DIV8, the VU treatment was stopped and cultures were maintained in normal medium until DIV21. We verified that at DIV21, E_GABA_ and GABA DF in CA1 pyramidal neurons in VU-treated slices were back to control levels (Fig. 6A,B), indicating that chloride levels were fully restored.

**Fig. 6.**
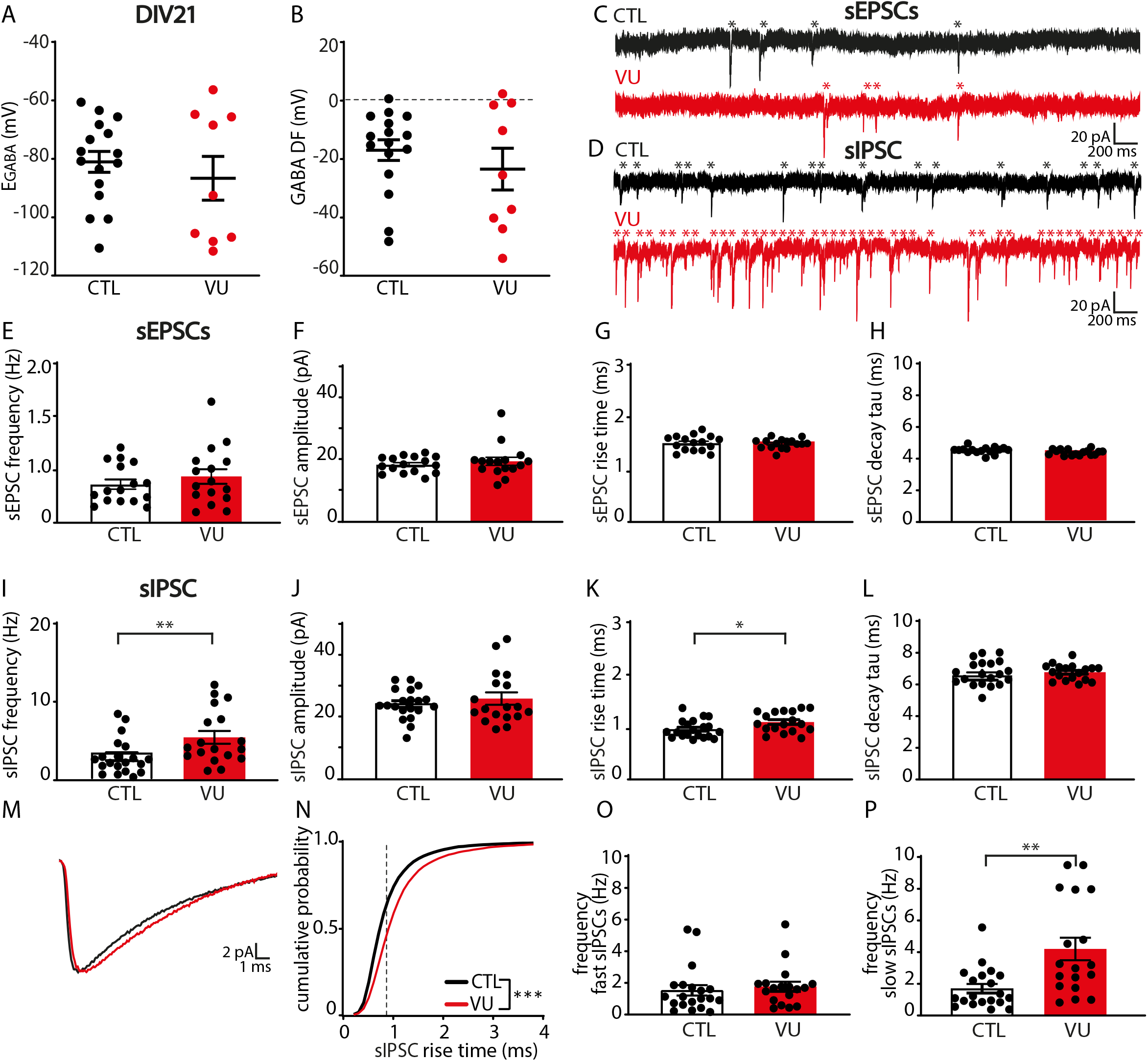
VU treatment does not affect excitatory transmission, but increases spontaneous inhibitory transmission at DIV21. A,B) GABA reversal potential (E_GABA_) (MW, p=0.56) and GABA Driving Force (GABA DF) (MW, p=0.76) recorded in CA1 pyramidal cells in control (CTL) and VU-treated cultures at DIV21, two weeks after cessation of treatment. Data from 9-16 cells, 3-10 slices and 3-10 mice per group. C, D) sEPSC and sIPSC recordings from CA1 pyramidal cells in CTL and VU-treated cultures at DIV21. sEPSCs and sIPSCs are indicated with asterisks. E-H) sEPSC frequency (UT, p=0.26), amplitude (MW, p=0.59), risetime (UT, p=0.80) and decay tau (UT, p=0.94) in CTL and VU-treated cultures at DIV21. Data from 16 cells, 9 slices and 9 mice per group. I-L) sIPSC frequency (MW, p=0.006), amplitude (UT, p=0.52), risetime (UT, p=0.026) and decay tau (UT, p=0.37) in CTL and VU-treated cultures at DIV21. M) Average sIPSC recorded in CA1 pyramidal cells in CTL and VU-treated cultures at DIV21. N) Cumulative distribution of sIPSC risetimes in control and VU-treated cultures at DIV21 (KS, p<0.0001). Dotted line indicates value used to split fast from slow rise time sIPSCs. O) Frequency of fast sIPSCs in CTL and VU-treated cultures at DIV21 (MW, p=0.20). P) Frequency of fast slow sIPSCs in CTL and VU-treated cultures at DIV21 (MW, p=0.003). Data in I-P from 18-20 cells, 11-12 slices and 8-9 mice per group.

We performed whole-cell recordings in CA1 pyramidal cells to record sEPSCs and sIPSCs (Fig. 6C,D). Similar to DIV9, we found no differences in sEPSCs between control and VU-treated slices at DIV21 (Fig. 6E-H). Surprisingly, we observed that sIPSC frequency in VU-treated slices was increased compared to control slices at DIV21 (Fig. 6I), whereas sIPSC amplitudes were not different (Fig. 6J). When we analyzed the kinetics of the individual sIPSCs we noticed that sIPSCs in VU-treated slices had slightly larger rise times compared to control (Fig. 6K), while decay times were not different (Fig. 6L). The IPSC rise time depends on the location of synapse where the current originates from, with somatic synapses generating currents with faster rise times while currents from dendritic synapses will be slower due to dendritic filtering (Rall, 1967; Bekkers and Clements, 1999; Wierenga and Wadman, 1999; Ruiter et al., 2020). When we split slow and fast sIPSCs (Ruiter et al., 2020), we observed that the increase was most prominent in slow sIPSCs (Fig. 6M-P), possibly reflecting an increase in IPSCs originating from dendritic synapses.

To assess whether the increase in inhibitory transmission after VU was activity dependent, we also measured mIPSCs in control and VU-treated slices at DIV21. We found that mIPSCs were not different in VU-treated and control slices (Fig. 7A-D), suggesting that VU treatment increased activity-dependent GABA release at DIV21. To assess possible changes in synaptic density, we determined the density and size of VGLUT and VGAT puncta in the CA1 area and we found no differences between control and VU-treated slices, both in sPyr and sRad (Fig. E-L). Spine density in CA1 pyramidal neurons was also not different in control and VU-treated slices at DIV21 (Fig. 7M,N). This indicates that excitatory and inhibitory synaptic density remained unaffected 14 days after the VU treatment and suggests that the observed increase in sIPSC frequency is due to an increased activity-dependent release from GABAergic terminals, possibly preferentially at dendritic inhibitory synapses.

**Fig. 7.**
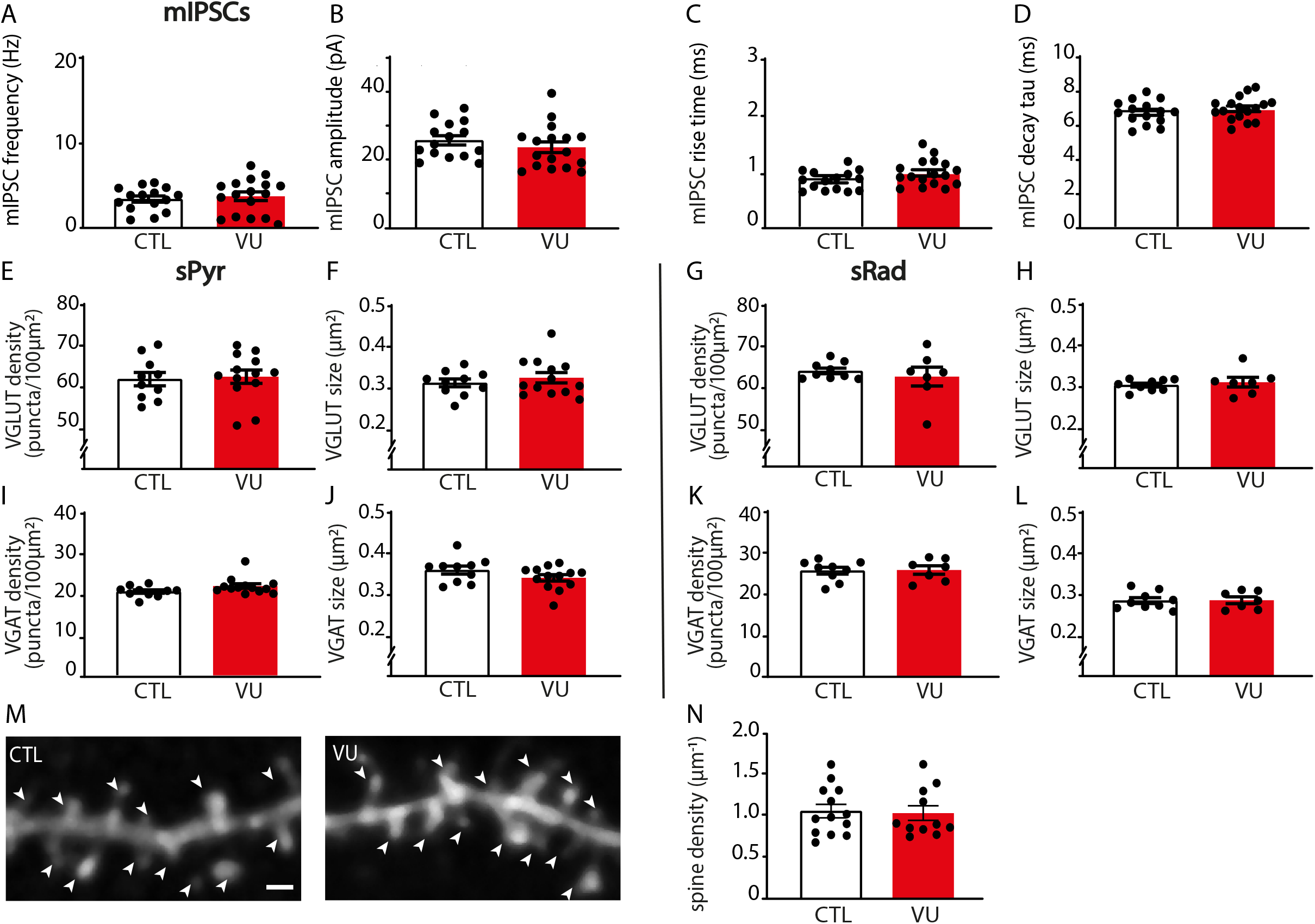
VU treatment does not affect mIPSCs, or excitatory and inhibitory synapses at DIV21. A-D) mIPSC frequency (UT, p=0.60), amplitude (UT, p=0.34), risetime (UT, p=0.077) and decay tau (UT, p=0.40) recorded from CA1 pyramidal cells in CTL and VU-treated cultures at DIV21. Data from 15-17 cells, 9-12 slices and 6-9 mice per group. E,F) Density (UT, p=0.80) and size (UT, p=0.61) of VGLUT puncta in the sPyr in CTL and VU-treated slices at DIV21. G,H) Density (MW, p=0.91) and size (MW, p=0.76) of VGLUT puncta in the stratum Radiatum (sRad). I,J) Density (MW, p=0.078) and size (UT, p=0.14) of VGAT puncta in the sPyr. K,L) Density (MW, p=0.76) and size (MW, p>0.99) of VGAT puncta in the sRad. Data in E-L from 10-13 slices and 6-7 mice per group. M) Maximal projection of confocal images of the apical dendrite of DIV21 CA1 pyramidal neurons in CTL and VU-treated cultures. Spines are indicated with arrowheads. Scale bar: 1 μm. N) Dendritic spine densities in control and VU-treated cultures at DIV21 (MW, p=0.77). Data from 10-11 cells, 8 slices and 7-8 mice per group.

Network discharges during sEPSC recordings were rare at DIV21 and not different between control and VU-treated slices (Fig. 8A-C). One week VU treatment did not affect firing properties (Fig. 8D,E) or resting membrane potential (RMP) (Fig. 8F) of CA1 pyramidal neurons at DIV21, but we observed a slight elevation of the action potential (AP) threshold in VU-treated slices at DIV21 (Fig. 8G). The relative AP threshold (defined as the difference between AP threshold and RMP, Fig. 8H) was not different VU-treated and control pyramidal cells. It is noteworthy that the AP threshold was not different when we recorded with high chloride internal solution, which resulted in a more negative AP threshold (DMSO -41.9 ± 3.3 mV; VU -42.9 ± 3.4 mV; UT, p=0.45; data not shown). A similar decrease in AP threshold after chloride loading was recently described in CA3 pyramidal neurons (Sørensen et al., 2017), but the mechanism remained unresolved.

**Fig. 8.**
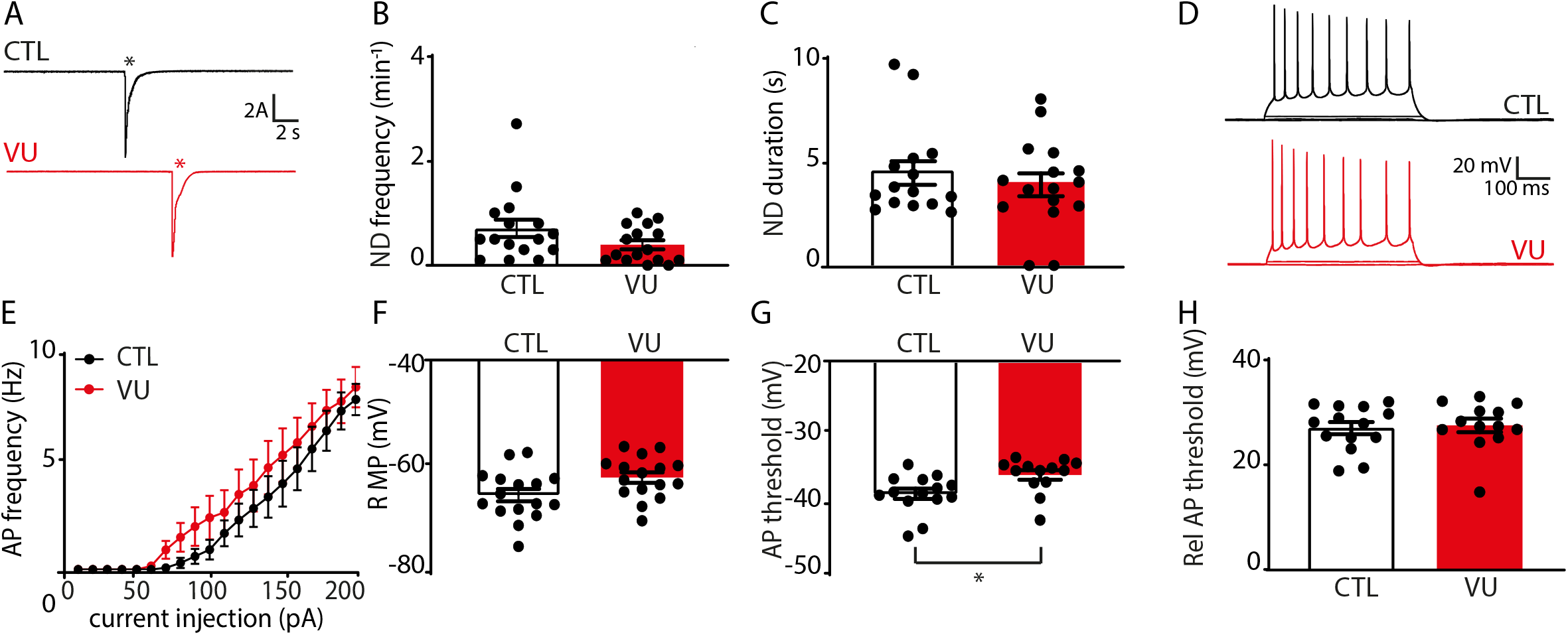
VU treatment increases the firing threshold of pyramidal neurons at DIV21. A) Whole-cell voltage clamp recordings from CA1 pyramidal cells in control (CTL) and VU-treated cultures at DIV21, in the presence of gabazine. Network discharges are indicated with asterisks. B, C) Network discharge (ND) frequency (MW, p=0.14) and duration (MW, p=0.20) in CTL and VU-treated cultures at DIV21. Data from 16 cells, 9 slices and 9 mice per group. D) Whole-cell current clamp recordings of action potentials after current injections in CA1 pyramidal neurons in CTL and VU-treated cultures at DIV21. E) Action potential firing rates in CTL and VU-treated cultures with increasing current injections at DIV21 (2W ANOVA, current injection: p<0.001, treatment: p=0.36). Data from 13-14 cells, 9 slices and 9 mice per group. F-H) Resting membrane potential (RMP) (UT, p=0.75), action potential (AP) threshold (MW, p=0.006), and relative action potential (Rel AP) threshold (MW, p=0.72) in CTL and VU-treated cultures at DIV21. Data from 13-16 cells, 9 slices and 9 mice per group.

### VU treatment does not affect chloride levels in interneurons at DIV9 or DIV21

Our data suggest that VU treatment specifically affected activity-dependent GABAergic transmission from interneurons to pyramidal neurons. We wondered if VU treatment affected chloride concentrations in inhibitory neurons in the same way as in pyramidal neurons. We therefore expressed the chloride sensor SClm (Grimley et al., 2013) specifically in GABAergic interneurons, using a Cre-dependent adeno associated virus (Boffi et al., 2018) in slices from VGAT-Cre mice. We measured SClm FRET ratios in cell bodies of GABAergic interneurons in sRad and sPyr (Fig. 9A). We observed that FRET values in interneurons were much lower compared to pyramidal cells at DIV9 (Fig. 2E), and clearly increased between DIV9 and DIV21 (Fig. 9B,C), reflecting a decrease of intracellular chloride levels with development. This was remarkable as we previously showed that chloride levels in pyramidal neurons are already mature at DIV9 and remain relatively stable during the this period (Herstel et al., 2022). When we normalized FRET ratios at DIV21 to DIV9 for interneurons and pyramidal neurons, the relative increase was 1.33 ± 0.03% in interneurons compared to 1.02 ± 0.03% in pyramidal neurons (UT, p<0.0001). This suggests that the GABA shift occurs later in inhibitory neurons compared to excitatory neurons in the CA1 area, which is in line with previous reports (Patenaude et al., 2005; Otsu et al., 2020). Interestingly, and in clear contrast to pyramidal neurons (Fig. 2G,F), VU treatment from DIV2 to DIV8 did not affect SClm FRET values in interneurons at DIV9 (Fig. 9B-E), suggesting that the contribution of KCC2 to chloride levels in interneurons was minimal during the treatment period. FRET ratios increased similarly to DIV21 in VU-treated and control slices (Fig. 9B-E). These results indicate that the one week VU treatment had a differential effect on intracellular chloride levels in CA1 pyramidal and interneurons, probably because of cell type-specific timing of chloride maturation.

**Fig. 9.**
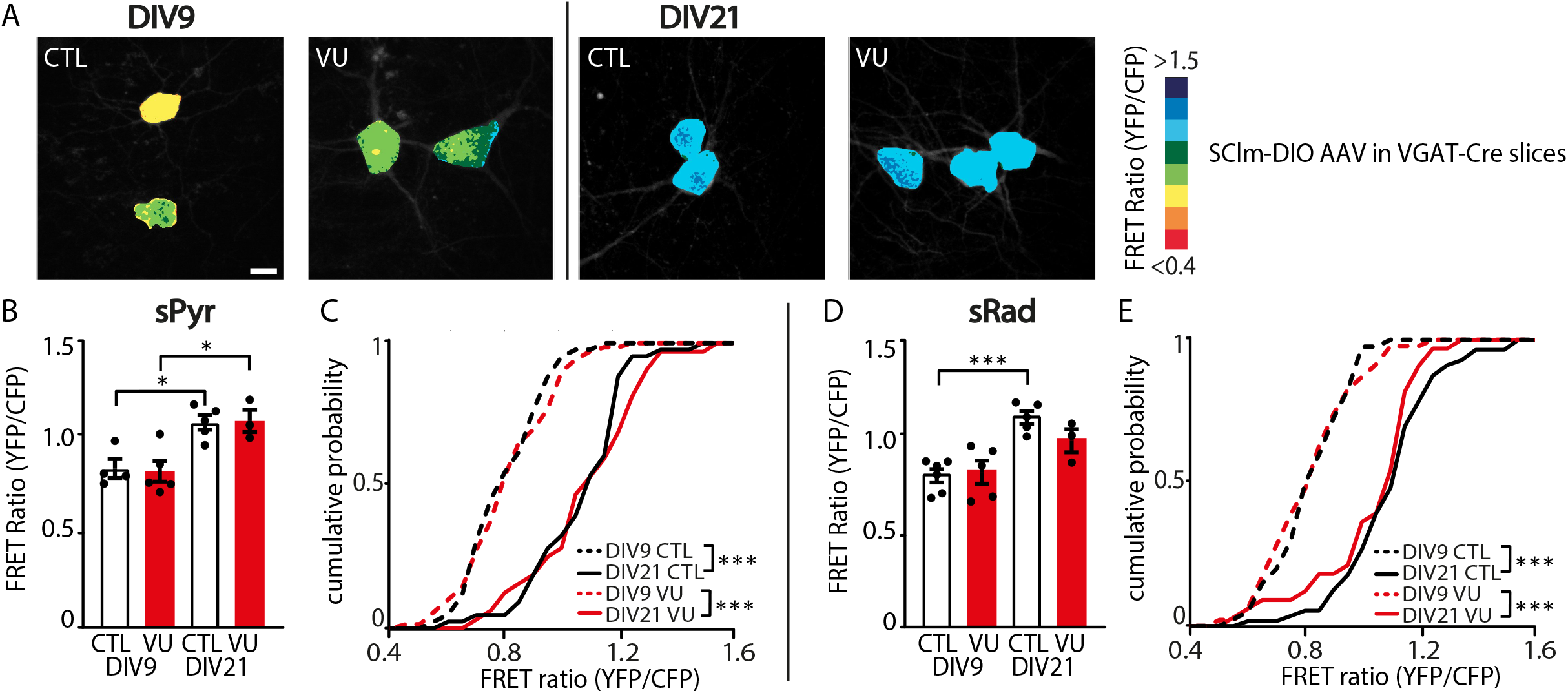
VU treatment does not change chloride concentrations in VGAT positive interneurons. A) Images of SClm FRET ratios in CA1 sRad GABAergic interneurons in control (CTL) and VU-treated slices from VGAT-Cre mice at DIV9 and 21. Scale bar: 10 μm. B) Average SClm FRET ratio in interneurons in the CA1 sPyr in CTL and VU-treated cultures at DIV9 and DIV21 (1W ANOVA, p=0.002; SMC: DIV9 DMSO versus DIV9 VU, p=0.99; DIV9 DMSO versus DIV21 DMSO, p=0.015; DIV9 VU versus DIV21 VU, p=0.015; DIV21 DMSO versus DIV21 VU, p=0.97). C) Cumulative distribution of FRET ratios in individual sPyr interneurons in CTL and VU-treated cultures (KS, p<0.0001). Data in B,C from 32-69 cells, 3-5 slices and 3 mice per group. D) Average SClm FRET ratio in interneurons in the CA1 sRad in CTL and VU-treated cultures at DIV9 and DIV21 (1W ANOVA, p=0.02; SMC: DIV9 DMSO versus DIV9 VU, p=0.77; DIV9 DMSO versus DIV21 DMSO, p=0.0007; DIV9 VU versus DIV21 VU, p=0.12; DIV21 DMSO versus DIV21 VU, p=0.23). An additional 1W ANOVA was performed to compare FRET ratios of individual cells in DIV9 VU versus DIV21 VU cultures and DIV21 DMSO versus DIV21 VU cultures (1W ANOVA, p<0.0001; SMC: DIV9 VU versus DIV21 VU, p<0.0001; DIV21 DMSO versus DIV21 VU, p=0.37). E) Cumulative distribution of FRET ratios in individual interneurons the CA1 sRad VU-treated (KS, p<0.0001) and VU-treated cultures (KS, p<0.0001). An additional KS was performed to compare the distribution of FRET ratios in CTL and VU-treated cultures at DIV21 (KS, p=0.63). Data in D,E from 33-56 cells, 3-6 slices and 2-3 mice per group.

### VU treatment results in elevated membrane potential in sRad interneurons at DIV21

As sIPSC frequency was increased in VU-treated slices at DIV21, while mIPSCs and VGAT puncta were unaffected, we wondered if release probability at GABAergic synapses was increased. We therefore assessed paired-pulse ratios (PPR) of evoked IPSCs in CA1 pyramidal cells after stimulation with an extracellular electrode in the sPyr or sRad (Fig. 10A), but PPR of IPSCs was similar in VU-treated and control slices (Fig. 10B,C). We also determined the coefficient of variation of the first IPSCs, another parameter that is dependent on release probability, but we did not find any difference between VU-treated and control slices (Fig. 10D,E). This suggests that IPSC release probability was not affected by VU treatment. We therefore wondered if the increase in sIPSC frequency in VU-treated slices could be due to an increased activity of GABAergic cells at DIV21. Based on our observation of a specific increase in sIPSC with slow rise times in VU-treated slices (Fig. 6M-P), we focused on dendritically targeting GABAergic cells in the sRad. We performed whole-cell patch clamp recordings from GFP-labeled interneurons in the sRad in slices from GAD65-GFP mice, which are mostly reelin-positive and preferentially target pyramidal dendrites (Wierenga et al., 2008, 2010). We found that sEPSCs (Fig. 10F-H), and general firing properties (Fig. 10I,J) of sRad interneurons were not different in VU-treated and control slices at DIV21. Intriguingly, we observed that in VU-treated slices resting membrane potential of the GFP-labeled interneurons was slightly elevated (Fig. 10K). The AP threshold was not significantly altered (Fig. 10L), but the relative action potential threshold was lower in VU-treated slices compared to control (Fig. 10M). Input resistance was not different (DMSO 271.1 ± 12.8 MΩ; VU 244.2 ± 13.2 MΩ; UT, p=0.15). Together, this suggests that sRad interneurons in VU-treated slices receive normal excitatory synaptic input at DIV21, but that less input current is required to reach their firing threshold. We propose that this subtle alteration in membrane excitability contributes to the elevated sIPSC frequency that we observed after VU treatment at DIV21.

**Fig. 10.**
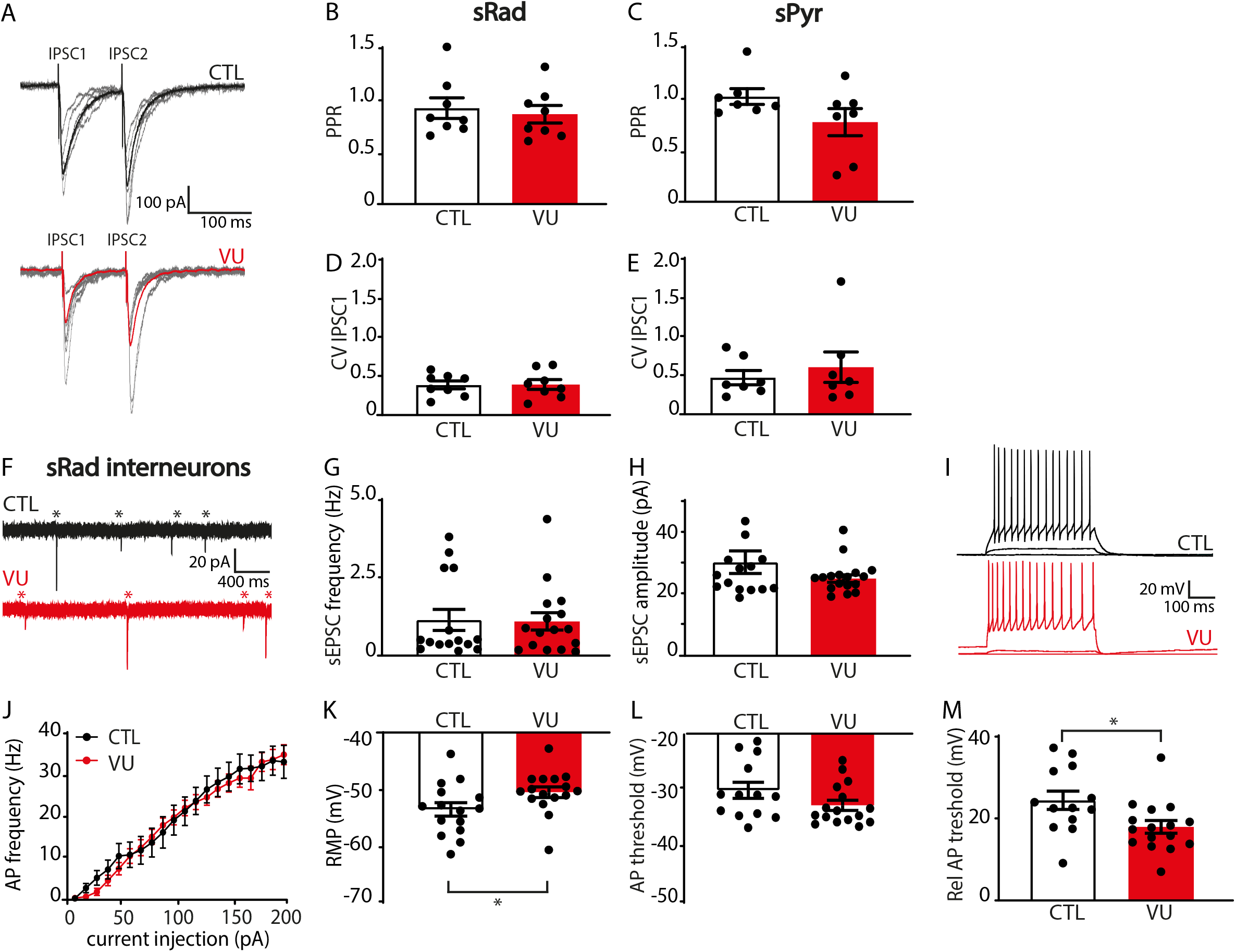
VU treatment results in elevated resting membrane potentials in sRad interneurons at DIV21. A) Whole-cell voltage clamp recordings of evoked IPSCs in control (CTL) and VU-treated cultures at DIV21. Stimulation electrode was placed in the sRad. Average evoked IPSCs are shown in bold, individual traces are shown in grey. B, C) Paired pulse ratios (PPR; IPSC2/IPSC1 amplitudes) evoked in sRad (MW, p=0.65) and sPyr (MW, p=0.21) in CTL and VU-treated cultures at DIV21. D, E) Coefficient of variation (CV) of the first IPSCs evoked in sRad (MW, p=0.96) and sPyr (MW, p=0.80) in CTL and VU-treated cultures at DIV21. Data in B-E from 7-8 cells, 4-6 slices and 3-5 mice per group. F) sEPSC recording in GFP-labeled interneurons in sRad from control (CTL) and VU-treated cultures from GAD65-GFP mice at DIV21. sEPSC are indicated with asterisks. G,H) sEPSC frequency (MW; p=0.85) and amplitude (MW; p=0.38) in these interneurons. Data from 15-16 cells, 11 slices and 10 mice per group. I) Whole-cell current clamp recordings of action potentials after current injections in sRad interneurons in CTL and VU-treated cultures at DIV21. J) Action potential firing rates in sRad interneurons in CTL and VU-treated cultures at DIV21 (2W ANOVA, current injection: p<0.001, treatment: p=0.42). K-M) Resting membrane potential (RMP) (MW, p=0.023), action potential (AP) threshold (UT, p=0.11), relative action potential (Rel AP) threshold (MW, p=0.012) in sRad interneurons in CTL and VU-treated cultures at DIV21. Data in J-M from 13-16 cells, 11 slices and 10 mice per group.

## Discussion

There is accumulating evidence that the developmental GABA shift is often delayed in NDD patients (Talos et al., 2012; Duarte et al., 2013; Merner et al., 2015; Tang et al., 2016; Ruffolo et al., 2018; Wang et al., 2021; Birey et al., 2022) and in NDD animal models (He et al., 2014; Tyzio et al., 2014; Deidda et al., 2015b; Banerjee et al., 2016; Corradini et al., 2017; Fernandez et al., 2018; Roux et al., 2018; Lozovaya et al., 2019; Bertoni et al., 2021). Here, we used organotypic hippocampal cultures to examine the consequences of a delayed GABA shift for the developing CA1 network using the specific KCC2 blocker VU0463271 (VU). VU treatment between DIV2 and DIV8 increased intracellular chloride levels without affecting chloride transporter expression levels, thereby effectively delaying the GABA shift in CA1 pyramidal neurons. We found that VU treatment did not have a direct effect on synaptic currents and firing properties of CA1 pyramidal cells at DIV9. However, at DIV21, when chloride levels were fully normalized, we observed a remarkable increase in sIPSC frequency compared to control slices, while synapse numbers remained unaffected. We found that firing thresholds in CA1 pyramidal neurons were slightly elevated while dendrite-targeting interneurons showed an elevated resting membrane potential and lower firing threshold at DIV21. Together, this shows that a delay in the postnatal GABA shift does not directly affect synaptic development, but rather leads to indirect, cell type-specific changes in membrane properties that may contribute to altered network activity at a later time point.

In this study we used organotypic hippocampal cultures as a model system to study postnatal development. We demonstrate that the GABA shift occurs in slice cultures around the same time as it does *in vivo* (Valeeva et al., 2016; Sulis Sato et al., 2017; Murata and Colonnese, 2020; Salmon et al., 2020; Herstel et al., 2022). As previously reported, synapses continue to develop in these cultured slices. Indeed, sEPSC frequency and density of VGLUT puncta and density of spines increased from DIV9 to DIV21 (De Simoni et al., 2003; Berry et al., 2012), while sEPSC amplitude decreased (De Simoni et al., 2003) and the frequency of sIPSCs and density of puncta VGAT remained mostly stable (De Simoni et al., 2003). VU treatment from DIV2 to DIV8 resulted in increased chloride levels in CA1 pyramidal cells at DIV9, that were comparable to untreated slices at DIV2. This indicates that the VU treatment effectively delayed chloride maturation and maintained depolarizing GABA signaling up to DIV9. Chloride maturation in interneurons occurs later compared to pyramidal cells and remained unaffected by the VU treatment. Importantly, VU treatment blocked KCC2 function without altering expression levels of the protein. This suggests that VU treatment altered intracellular chloride levels in pyramidal cells without interfering with the structural role of KCC2 proteins in spines (Li et al., 2007; Gauvain et al., 2011; Fiumelli et al., 2013; Chevy et al., 2015; Puskarjov et al., 2017; Awad et al., 2018; Kesaf et al., 2020). Our experimental design therefore allowed for selectively testing how intracellular chloride concentrations influence the developing CA1 network.

The first key finding from this study is that maintaining depolarizing GABA signaling until DIV9 did not have any direct effects on excitatory and inhibitory synaptic structure or function. Previous groundbreaking research has demonstrated that excitatory GABAergic signaling during prenatal and perinatal development promotes synapse formation and maturation (Leinekugel et al., 1997; Akerman and Cline, 2006; Nakanishi et al., 2007; Wang and Kriegstein, 2008, 2011; van Rheede et al., 2015; Oh et al., 2016). These studies were performed before P8, or when the GABA shift was accelerated. Our slices are made from P6-7 pups, when GABA signaling is still depolarizing. Here we show that prolonging the period of depolarizing GABA signaling during the second postnatal week does no longer affect synapse formation. We did not observe any difference in synapse number between control and VU-treated slices, at DIV9 nor at DIV21. This suggests that the positive effect of depolarizing GABA signaling is restricted from embryonic development up to the first postnatal week. In our slices, even when GABA signaling was kept depolarizing, GABA signaling already had an inhibitory action on cellular and network activity. Indeed, we found that although E_GABA_ in VU-treated slices was depolarized relative to RMP at DIV9 (Fig. 2B), it remained well below AP threshold (Fig. 3N). As a result, GABA signaling was inhibitory in both control and VU-treated slices (Fig. 5). The effect of GABA signaling under physiological conditions does not only depend on the relation between E_GABA_ and the AP threshold, but also on the spatiotemporal distribution of glutamatergic and GABAergic inputs (Staley and Mody, 1992; Gao et al., 1998; Morita et al., 2006; Le Magueresse and Monyer, 2013; Kilb, 2021). With more (excitatory) synaptic activity, shunting inhibition will become more prominent (Woodruff et al., 2011; Branchereau et al., 2016). This means that the precise consequences of alterations in the postnatal GABA shift depend on the effect on local network activity (Wang and Kriegstein, 2011; Seja et al., 2012; Deidda et al., 2015a; Pisella et al., 2019). Our results are in line with previous reports which argue that depolarizing GABAergic signaling is inhibitory in hippocampal slices from approximately P6-9 (Khalilov et al., 1997; Valeeva et al., 2016; Salmon et al., 2020). Also in the hippocampus of living mice, depolarizing GABA was found to inhibit activity in the hippocampus from P7 onwards (Valeeva et al., 2010; Murata and Colonnese, 2020). Our results therefore support the notion that depolarizing but inhibitory GABA signaling has limited contribution to postnatal synapse development in the hippocampus.

The second key finding from this study is that subtle, and cell type specific, alterations in membrane properties were observed two weeks after the VU treatment had ended. We found that CA1 pyramidal cells had a slightly increased firing threshold, and interneurons in the sRad showed a slightly elevated membrane potential, which reduced the amount of depolarization required to fire an AP. We speculate that together this will modify spontaneous network activity toward an increased inhibitory tone and may explain the increased sIPSC frequency that we observed in VU-treated slices (Fig. 6I). Interestingly, pharmacological inhibition of NKCC1 from P3 to P8 (resulting in decreased intracellular chloride levels) has been reported to transiently decrease inhibitory transmission in the visual cortex several weeks later (at P35) (Deidda et al., 2015a), suggesting that developing GABAergic transmission may be particularly sensitive to intracellular chloride levels.

It remains unclear how blocking KCC2 from DIV2 to DIV8 can alter membrane properties of neurons two weeks later. Our study adds to an increasing number of studies that demonstrate that intracellular chloride levels can modify ion channels, and therefore membrane excitability, in often unpredictable ways (Huang et al., 2012; Seja et al., 2012; Goutierre et al., 2019; Sinha et al., 2022). Most notably, it was shown that membrane levels of specific potassium channels are regulated via KCC2 (Goutierre et al., 2019) and the chloride-dependent kinase WNK3 (Sinha et al., 2022), and it will be important to further examine the role of various chloride channels in membrane excitability (Jentsch, 2016; Jentsch and Pusch, 2018; Akita and Fukuda, 2020). Interestingly, effects seem to strongly depend on cell type (Seja et al., 2012) and timing of the chloride manipulation (Lim et al., 2021; Sinha et al., 2022). The changes in excitability that we observed at DIV21 may therefore reflect a complex sequence of subtle adaptations of ion channel distribution that is specific per cell type.

The present work shows that delaying the postnatal GABA shift by one week has no direct effects on synaptic development. Instead, we found evidence for indirect, cell type-specific changes in membrane properties, possibly via chloride-dependent regulation of ion channels, which may have long-term consequences for network activity and brain function. Our data call for careful assessment of alterations in cellular excitability in NDDs.

## 11. Acknowledgements

This research was supported by a TOP grant from ZonMW (#91216021) and by the Nederlandse Organisatie voor Wetenschappelijk Onderzoek (NWO; #OCENW.KLEIN.150). We thank Prof. Thomas Kuner for providing the Cre-dependent SCLM construct (Boffi et al., 2018). We thank Prof. Kevin Staley for providing the SuperClomeleon^lox/-^ mouse line, Dr. Stefan Berger for the CamKIIα^Cre/-^ mice and Dr. Henk Karst for sharing these mice with us. We thank Dunya Selemangel for help with SuperClomeleon data analysis and René van Dorland for the AAV production and excellent technical support.

